# Albumin-fused interleukin-10 prevents experimental autoimmune encephalomyelitis by promoting immunoregulation in the secondary lymphoid organs

**DOI:** 10.64898/2026.04.10.717764

**Authors:** Erica Budina, Joseph W. Reda, Kirsten C. Refvik, Julia Luehr, Brendan T. Berg, Hye-Rin Chun, Taryn N. Beckman, Ani Solanki, Mindy Nguyen, Sofia N. Reda, Colleen R. Foley, Ivan Vuong, Abigail L. Lauterbach, Kevin Hultgren, Suzana Gomes, Jun Ishihara, Lisa R. Volpatti, Jeffrey A. Hubbell

## Abstract

Interleukin-10 (IL-10) is an immunoregulatory cytokine that suppresses pro-inflammatory cytokine production, myeloid cell antigen presentation, and T cell activation. However, the therapeutic use of recombinant IL-10 has been limited by its short plasma half-life and limited tissue retention, including in the secondary lymphoid organs (SLOs), key sites of autoreactive T cell priming in autoimmune disease. We previously engineered a fusion of serum albumin and IL-10 (SA-IL-10) with extended plasma half-life and enhanced accumulation in the SLOs. Here, we combine analysis of human single-cell transcriptomic data with studies in a murine experimental autoimmune encephalomyelitis (EAE) model to investigate how sustained IL-10 exposure in the SLOs modulates immune responses during neuroinflammation. Single-cell RNA-sequencing analysis of treatment-naïve multiple sclerosis (MS) patients revealed a significantly elevated percentage of regulatory CD4^+^ T cells in the cerebrospinal fluid expressing *IL10RA*, the ligand-binding chain of the IL-10 receptor, whereas *IL10* transcript detection was low and not different in MS compared to controls. Prophylactic subcutaneous administration of SA-IL-10 prevented EAE development and progression with superior efficacy to wild type IL-10 and comparable protection to fingolimod. In the SLOs, SA-IL-10 reduced antigen-specific T_H_17 cells, M1-like macrophages, activated dendritic cells, and antigen-induced pro-inflammatory cytokine production, while expanding M2-like macrophages and elevating the frequencies of multiple checkpoint markers on T_H_2 cells. SA-IL-10 also reduced immune cell infiltration in the spinal cord. Repeated SA-IL-10 administration was well tolerated in EAE, with plasma biochemistry parameters comparable to controls. Together, these findings demonstrate that increasing IL-10 exposure in the SLOs suppresses neuroinflammation by promoting immunoregulation.

## Introduction

Interleukin-10 (IL-10) is an immunoregulatory cytokine secreted by myeloid and lymphoid cells, including macrophages, monocytes, dendritic cells, neutrophils, mast cells, eosinophils, natural killer cells, CD4^+^ and CD8^+^ T cell subsets, and B cells.^1,2^ By engaging the IL-10Rα:IL-10Rβ signaling complex, IL-10 suppresses pro-inflammatory cytokine production, reduces antigen presentation by myeloid cells, promotes M2 macrophage polarization, and inhibits T cell activation.^3^ IL-10 suppresses T_H_1 and T_H_17-associated responses by downregulating antigen presentation by activated dendritic cells (DCs) and macrophages,^1,4^ reducing DC migration to the lymph nodes (LNs),^5,6^ and inhibiting CD4^+^ T cell proliferation and effector cytokine secretion.^7,8^ Through these mechanisms, IL-10 contributes to immune homeostasis.^9^

Despite its potent immunoregulatory functions, efforts to harness recombinant IL-10 as a therapy have been limited by its unfavorable pharmacokinetic properties, including its short plasma half-life (the mean terminal-phase half-life of recombinant human IL-10 is 2.7 to 4.5 hr) and limited tissue persistence following systemic administration.^3,10,11^ Wild type (WT) IL-10 has a molecular weight below the threshold for renal filtration and lacks recycling mechanisms, resulting in rapid renal clearance and proteolytic degradation in the bloodstream.^12^ Consequently, IL-10 exhibits limited systemic persistence and poor accumulation in the secondary lymphoid organs (SLOs), critical sites of antigen presentation, immune cell activation, and autoreactive T cell priming in autoimmune diseases.^12^

To address these limitations, we previously engineered a fusion protein consisting of IL-10 recombinantly linked to serum albumin (SA),^11^ an abundant endogenous plasma protein with a long circulating plasma half-life,^13^ and characterized the pharmacokinetics and tissue localization of SA-IL-10 following intravenous administration.^11^ Compared to WT IL-10, SA-IL-10 exhibited prolonged half-life, yielded approximately 5- to 18-fold greater IL-10 exposure by area-under-the-curve in the popliteal, mesenteric, and cervical LNs, and remained detectable in the popliteal LNs for up to 72 hr.^11^ Imaging of LNs demonstrated that SA-IL-10 accumulated within the LN parenchyma and localized in proximity to high endothelial venules, suggesting a route of LN entry other than the afferent lymphatics.^11^ We further showed that SA-IL-10 treatment was more effective than WT IL-10 in both the passive collagen antibody-induced arthritis and active collagen-induced arthritis models, with efficacy comparable to anti-TNF*-α* antibody treatment, supporting the hypothesis that sustained cytokine exposure in the SLOs enhances the therapeutic activity of recombinant IL-10.^11^

Multiple sclerosis (MS) is a chronic, autoimmune, demyelinating, and neurodegenerative disease where immune dysregulation contributes to damage of the central nervous system (CNS).^14^ MS pathology is characterized by the activation and migration of autoreactive lymphocytes and innate immune cells into the CNS, contributing to demyelination, glial activation, axonal injury, and ultimately neurodegeneration.^14,15^ Although current MS therapies reduce relapse frequency and manage acute symptoms, many require long-term administration, broadly suppress the immune system, and display limited efficacy in halting progressive forms of the disease.^14^

MS arises from both the expansion of pathogenic immune cells and impaired immunoregulatory networks, resulting in defective suppressor cell function and dysregulated immune responses.^16,17^ Peripheral blood mononuclear cells from MS patients exhibit reduced *IL10* mRNA levels and elevated pro-inflammatory cytokine production compared to healthy controls.^18,19^ Regulatory T cells (Tregs) from MS patients display impaired suppressive function,^19,20^ while inflammatory monocytes and dendritic cells are expanded and exhibit increased activation marker expression and pro-inflammatory cytokine production.^21–23^ Several FDA-approved MS therapies, including glatiramer acetate,^24–26^ interferon-β,^27–30^ and fingolimod,^31,32^ have been shown to increase IL-10 production or IL-10 receptor signaling across multiple immune cell types and immune compartments, including the blood^33^ and cerebrospinal fluid (CSF).^34,35^

Despite the central role of IL-10-IL10R signaling in immunoregulation, administration of recombinant IL-10 has shown variable efficacy in experimental autoimmune encephalomyelitis (EAE), a preclinical model of MS.^36–39^ Studies in IL-10-deficient (IL-10^-/-^) C57BL/6 mice have shown that the loss of IL-10 exacerbates EAE,^40^ while transgenic expression of human IL-10 under the control of a class II MHC promoter in SJL/J x BALB/cAnN F1 mice confers resistance to active EAE development.^38^ Recombinant IL-10 administration has yielded divergent outcomes depending on species, strain, and method of EAE induction: disease was suppressed in Lewis rats with MOG_35-55_/CFA-induced EAE when treatment was administered either at the time of immunization or after disease onset,^36,37^ but not in SJL mice with adoptive transfer-induced acute EAE.^39^ These different outcomes may result from a combination of disease-specific factors and limited *in vivo* exposure of recombinant IL-10.

Here, we reanalyzed a published single-cell RNA sequencing (scRNA-seq) dataset comparing treatment-naïve MS patients to control individuals diagnosed with idiopathic intracranial hypertension (IIH), a non-inflammatory disorder of raised intracranial pressure,^41^ to examine IL-10 receptor and cytokine gene expression in human neuroinflammatory disease. This analysis revealed significantly higher percentages of regulatory CD4^+^ T cells expressing *IL10RA*, the ligand-binding chain of the IL-10 receptor,^3^ in the CSF of MS patients relative to IIH controls, whereas *IL10* transcript detection in immune cells was not measurably different between these populations. These data suggest regulatory CD4^+^ T cell populations in the CSF of MS patients retain the capacity to respond to IL-10, motivating the investigation of an exogenous IL-10 supplementation strategy. We then used EAE, a murine model of neuroinflammation, to investigate how sustained IL-10 exposure in the SLOs modulates immune responses during neuroinflammation. We showed that subcutaneously administered SA-IL-10 prevented the development and progression of EAE, with superior efficacy to WT IL-10 and comparable efficacy to fingolimod (FTY720), an FDA-approved MS drug. In the SLOs, SA-IL-10 administration reduced pathogenic, antigen-specific T_H_17 cells, M1-like macrophages, activated DCs and antigen-induced pro-inflammatory cytokine production, while expanding immunoregulatory M2-like macrophages and increasing the frequency of checkpoint markers (CTLA-4, PD-1, TIGIT, ICOS) on T_H_2 cells. SA-IL-10 treatment also reduced macrophage, DC, and CD4^+^ T cell infiltration in the spinal cord. Repeated SA-IL-10 administration was well tolerated, as treated EAE mice gained significantly more body weight over the course of treatment compared to PBS- and WT-IL-10 treated controls, and exhibited plasma biochemistry parameters comparable to control animals at study endpoint. These findings demonstrate that increasing IL-10 exposure in the SLOs suppresses neuroinflammation by promoting immunoregulation.

## Results

### Immune cells in the cerebrospinal fluid of treatment-naïve multiple sclerosis patients exhibit increased frequency of IL10RA expression in Treg CD4^+^ T cells

To characterize IL-10 pathway gene expression in MS, we re-analyzed published human scRNA-seq data from peripheral blood mononuclear cells (PBMCs) and CSF immune cells isolated from treatment-naïve MS patients and control individuals with IIH^41^ **(Figure 1a-b)**. Immune cell populations were clustered **(Supp. Figure 1a-b)** and annotated at the cluster level against the Monaco Immune Reference Dataset **(Supp. Figure 1c-f)**.^42^ We applied a confidence threshold of 0.6 and designated clusters that scored below this threshold for all Monaco broad labels as “Unlabeled” **(Supp. Figure 1c-f)**. Using this approach, we identified multiple cell types, including non-Treg CD4^+^ T cells, Treg CD4^+^ T cells, CD8^+^ T cells, B cells, NK cells, monocytes, DCs, and γδ T cells in each tissue, as well as a small population of progenitors in the blood **(Figure 1a-b)**.

**Figure 1.**
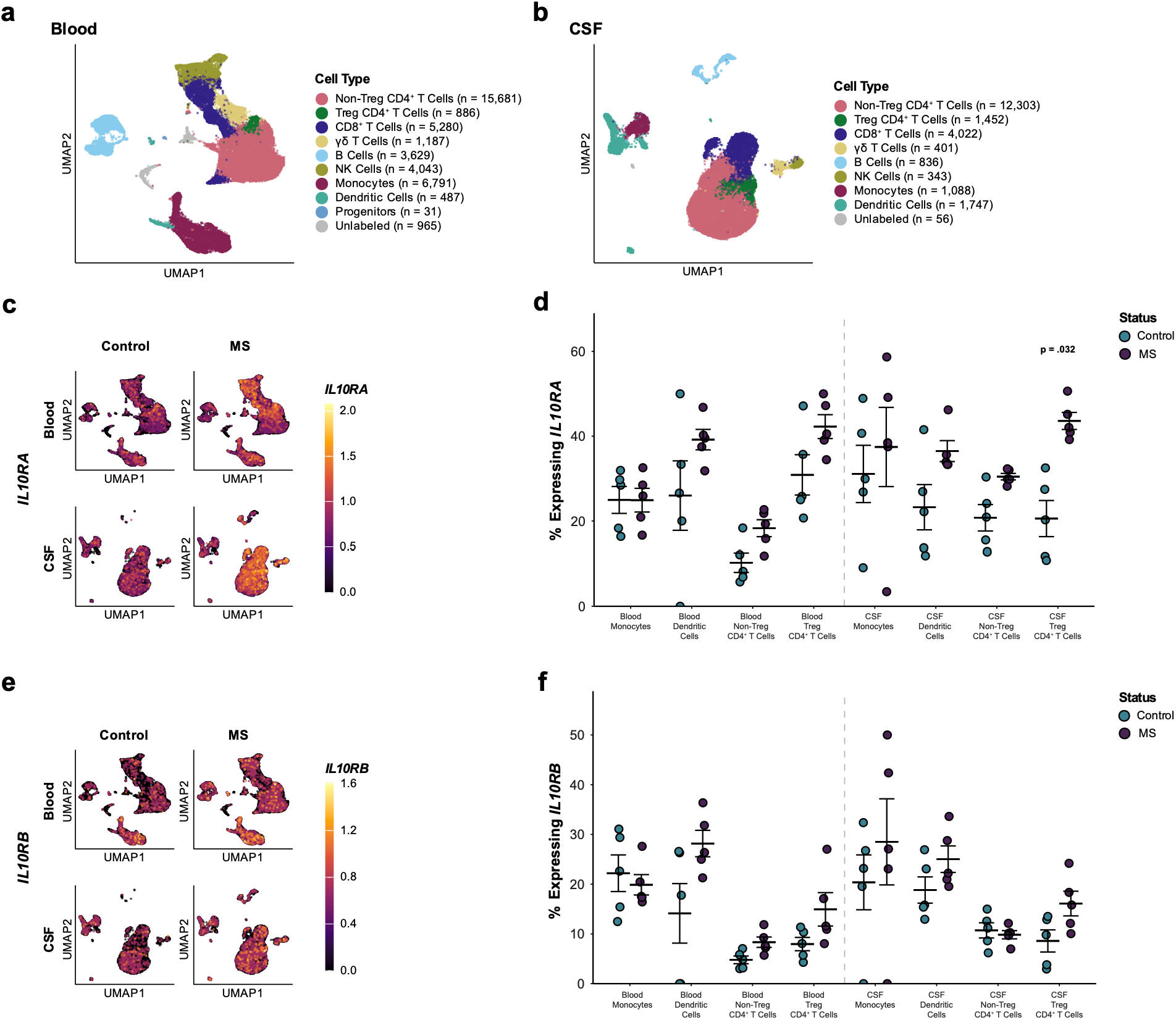
Treg CD4^+^ T cells in cerebrospinal fluid from MS patients exhibit increased *IL10RA* transcript expression. **a-b.** Uniform Manifold Approximation and Projection (UMAP) visualization of immune cell lineages in **(a)** blood and **(b)** cerebrospinal fluid (CSF), annotated at the cluster level. **c.** UMAP feature plots of *IL10RA* gene expression in blood- and CSF-derived immune cells from multiple sclerosis (MS) patients and control patients with idiopathic intracranial hypertension (IIH). **d.** Percentage of monocytes, dendritic cells, non-Treg CD4^+^ T cells, and Treg CD4^+^ T cells expressing the *IL10RA* gene in the blood and CSF. **e.** Feature plots of *IL10RB* gene expression in blood- and CSF-derived immune cells from MS patients and IIH controls. **f.** Percentage of monocytes, dendritic cells, non-Treg CD4^+^ T cells, and Treg CD4^+^ T cells expressing the *IL10RB* gene in the blood and CSF. Each data point in **(d)** and **(f)** represents the percentage of cells expressing *IL10RA* and *IL10RB*, respectively, per patient. Crossbars and whiskers represent mean ± s.e.m. Single-cell RNA sequencing data were obtained from dataset GSE138266 (MS patients, *n* = 5; IIH control patients, *n* = 5).^41^ Cell-type annotation was performed at the cluster level against the Monaco Immune Reference Dataset,^42^ with a confidence threshold of 0.6; clusters that scored below this threshold for all Monaco broad labels were designated “Unlabeled.” Statistical comparisons were performed using Wilcoxon rank-sum tests with Holm-Bonferroni correction for multiple comparisons within each gene and tissue; adjusted P values are displayed on the graphs for comparisons reaching P < 0.05.

To validate these assignments, we examined the expression of canonical lineage markers across all annotated populations in the blood and CSF of patients with MS and IIH controls **(Supp. Figure 2)**. Each annotated population preferentially expressed its corresponding canonical lineage markers **(Supp. Figure 2)**. The lineage markers expressed by each cell type were consistent between both blood and CSF, which were annotated independently **(Supp. Figure 2)**.

Subsequent analyses focused on monocytes, DCs, non-Treg CD4^+^ T cells, and Tregs **(Figure 1, Supp. Figure 3)**, selected based on their relevance to IL-10 signaling in neuroinflammation.^3^ While feature plot visualization showed widespread *IL10RA* and *IL10RB* expression in multiple immune populations (**Figure 1c** and **Figure 1e**), *IL10* was detected in relatively few cells **(Supp. Figure 3a-b)**. For each patient, we then quantified the percentage of cells within each population expressing *IL10RA*, *IL10RB,* and *IL10*, and compared these patient-level percentages between patients with MS and IIH controls **(Figure 1c-f, Supp. Figure 3b, Supp. Table 1)**.

In the CSF, the percentage of cells expressing *IL10RA*, the ligand-binding chain of the IL-10 receptor, was significantly higher among Treg CD4^+^ T cells in patients with MS than in IIH controls (Hedges’ *g* = 2.79; adjusted *P* value = 0.032) **(Figure 1c-d, Supp. Figure 3d, Supp. Table 1)**. The percentage of *IL10RA*-expressing CSF non-Treg CD4^+^ T cells (Hedges’ *g* = 1.73; adjusted *P* value = 0.167) and DCs (Hedges’ *g* = 1.29; adjusted *P* value = 0.187) in MS patients were also associated with large positive estimated effect sizes relative to IIH controls, but the differences were not statistically significant after correction for multiple comparisons **(Figure 1c-d, Supp. Table 1)**. Similarly, the percentage of *IL10RA*-expressing non-Treg CD4^+^ T cells in the blood of MS patients was associated with a large positive estimated effect size (Hedges’ *g* = 1.51) relative to IIH controls, but the difference was not statistically significant after correction for multiple comparisons (adjusted *P* value = 0.222) **(Figure 1c-d, Supp. Figure 3c, Supp. Table 1)**.

We also observed large positive estimated effect sizes for the percentage of *IL10RB*-expressing cells in MS patients relative to IIH controls. These populations included Treg CD4^+^ T cells (Hedges’ *g* = 1.29; adjusted *P* value = 0.381) and DCs (Hedges’ *g* = 0.94; adjusted *P* value = 0.667) in the CSF, as well as non-Treg CD4^+^ T cells (Hedges’ *g* = 1.56; adjusted *P* value = 0.127), Treg CD4^+^ T cells (Hedges’ *g* = 1.10; adjusted *P* value = 0.286), and DCs (Hedges’ *g* = 1.22; adjusted *P* value = 0.286) in the blood **(Figure 1e-f, Supp. Table 1)**. However, these comparisons were not statistically significant after correction for multiple comparisons **(Figure 1e-f, Supp. Table 1)**.

In the blood, *IL10* was detected in a low percentage of cells in the monocyte, DC, non-Treg CD4^+^ T cell, and Treg compartments **(Supp. Figure 3a-b)**. In the CSF, 4 out of 5 patients in each group demonstrated a higher percentage of *IL10* gene expression in monocytes than in DCs or non-Treg and Treg CD4^+^ T cells; however, the percentage of *IL10*-expressing cells did not differ significantly between patients with MS and IIH controls within any population **(Supp. Figure 3a-b, Supp. Table 1)**. Overall, these data show a selective increase in the percentage of Treg CD4^+^ T cells expressing *IL10RA* in treatment-naïve MS patients.

### Subcutaneous SA-IL-10 administration prevents the development and progression of MOG_35-55_-induced EAE

SA-IL-10 was generated by recombinantly fusing murine SA to murine IL-10 using a (GGGS)_2_ flexible linker, as described previously **(Supp. Table 2)**.^11,43^ SA-IL-10 was expressed in HEK 293-F cells and purified via affinity chromatography followed by size-exclusion chromatography, as described previously.^11,43^ We sought to evaluate the effect of prophylactic subcutaneous SA-IL-10 administration on EAE onset and progression. To this end, we induced active EAE in C57BL/6 mice by immunization with myelin oligodendrocyte glycoprotein (MOG)_35-55_ antigen emulsified in Complete Freund’s Adjuvant (CFA), followed by pertussis toxin to increase blood-brain barrier permeability.^44^

To identify a dose of SA-IL-10 capable of preventing disease onset, we administered PBS; 10 μg WT IL-10; or 2 μg, 10 μg, or 20 μg SA-IL-10 (all equimolar with respect to IL-10) subcutaneously every other day beginning on day 8 after disease induction **(Figure 2a)**. We selected subcutaneous administration due to its clinical convenience and potential to enable patient self-administration.^45^ Mice that received PBS, 2 μg SA-IL-10, or 10 μg WT IL-10 developed severe hindlimb paralysis by day 16, as indicated by the elevated EAE clinical scores and an increased percentage of mice with disease score *≥* 1 **(Figure 2b-c, Supp. Figure 4)**. In contrast, treatment with 10 μg or 20 μg SA-IL-10 (both equimolar with respect to IL-10) prevented disease development in nearly all mice **(Figure 2b-c, Supp. Figure 4)**.

**Figure 2.**
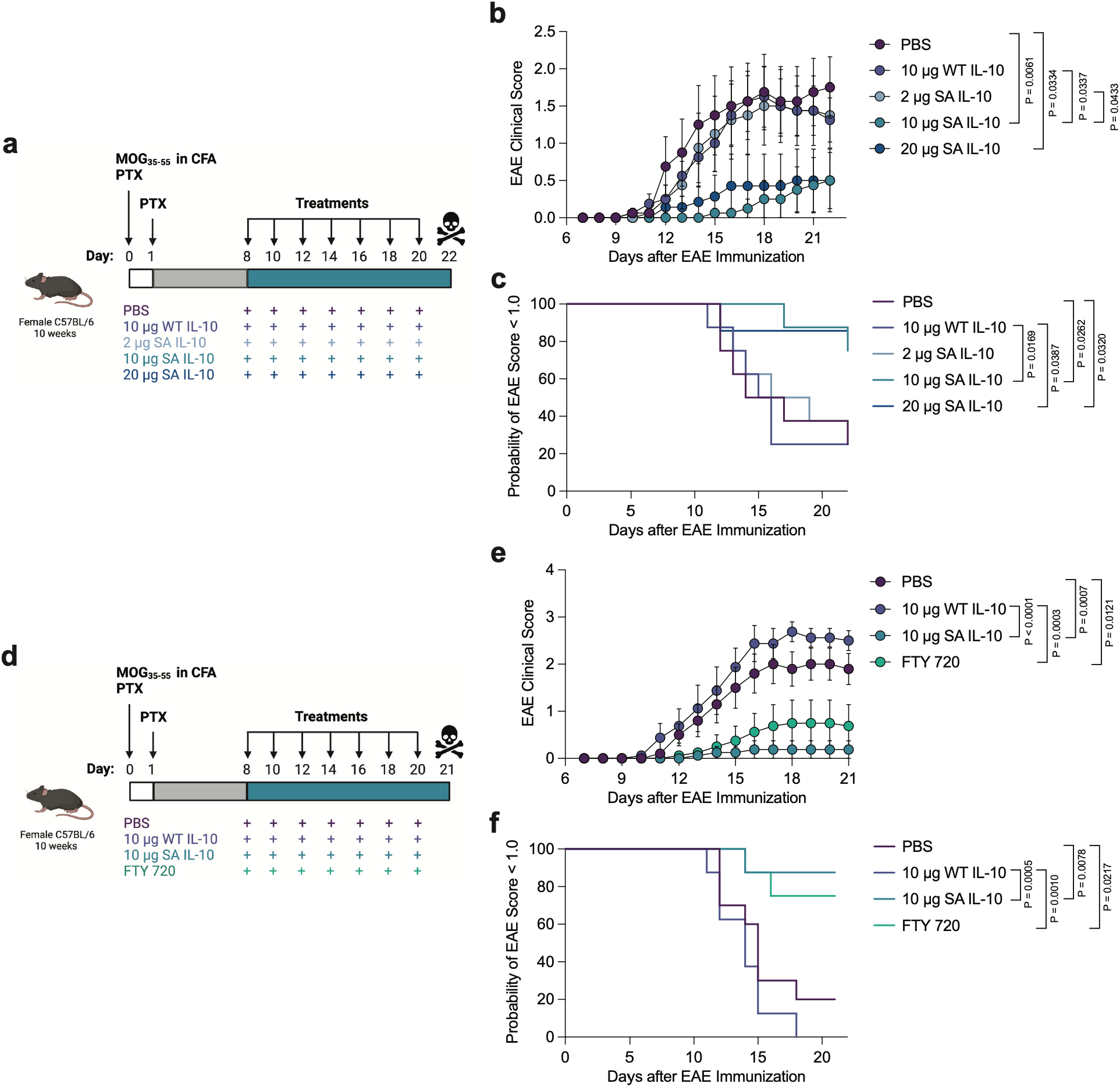
Prophylactic SA-IL-10 administration prevents the development and progression of experimental autoimmune encephalomyelitis. **a.** Overview of experimental design. Experimental autoimmune encephalomyelitis (EAE) was induced in C57BL/6 mice by s.c. immunization with MOG_35-55_/CFA followed by i.p. injections of PTX on days 0 and 1. Mice were prophylactically administered s.c. PBS, 10 μg WT IL-10, 2 μg SA-IL-10, 10 μg SA-IL-10, 20 μg SA-IL-10 (all molar e.q. to WT IL-10) every other day from day 8 to day 20 (*n* = 8 mice/group). Clinical scores were recorded daily from day 7 through the study endpoint. **b.** Disease progression. **c.** Disease incidence, indicated by the probability of disease score < 1.0. **d.** Overview of experimental design. EAE was induced as described in **(a)**. Mice were prophylactically administered s.c. PBS, 10 μg WT IL-10, 10 μg SA-IL-10 (all equimolar with respect to IL-10) every other day from day 8 to day 20, or i.g. FTY720 (1 mg/kg) daily from day 8 (*n* = 8-10 mice/group). **e.** Disease progression. **f.** Disease incidence. Data represent means ± s.e.m. Statistical analysis on disease score AUC (from d8 to d22 in **(b)** and d8 to d21 in **(e)**) was performed using one-way ANOVA with Tukey’s multiple-comparisons test, and on disease incidence in **(c, f)** using Log-rank (Mantel-Cox) test comparing every two groups. Diagrams in **(a, d)** were created using BioRender.com.

After identifying an optimal SA-IL-10 dose, we compared the efficacy of equimolar SA-IL-10 to WT IL-10, administered subcutaneously every other day beginning on day 8, and to FTY720, an FDA-approved MS drug,^46^ administered by daily oral gavage beginning on day 8 **(Figure 2d)**. We hypothesized that SA-IL-10’s extended bioavailability and prolonged exposure in the SLOs would enable improved disease prevention compared to unmodified WT IL-10. SA-IL-10 was significantly more effective than WT IL-10 at preventing EAE development and showed comparable efficacy to FTY720 **(Figure 2e-f, Supp. Figure 5)**. After long-term monitoring, only 1 out of the 8 SA-IL-10-treated mice exhibited disease symptoms, whereas 8 out of the 8 WT IL-10-treated mice and 2 out of the 8 FTY720-treated mice developed EAE by day 21, the experimental endpoint **(Figure 2e-f, Supp. Figure 5)**. These results suggest that albumin fusion to IL-10 enhances the prophylactic efficacy of IL-10 and prevents the onset and development of MOG_35-55_-induced EAE.

### SA-IL-10 reduces the activation of pro-inflammatory myeloid cells in the SC-dLNs of EAE-bearing mice

As the SLOs are critical sites of antigen presentation and myeloid-mediated T cell priming during EAE,^44^ we examined the impact of prophylactic SA-IL-10 treatment on myeloid cell populations in the cervical and iliac LNs (spinal cord draining lymph nodes, SC-dLNs) of EAE-bearing mice. For this purpose, we isolated cells from the SC-dLNs and performed immunophenotyping **(Figure 3a)**. SA-IL-10 administration significantly increased the percentage of CD11b^+^ F4/80^+^ macrophages in the SC-dLNs compared to PBS, WT IL-10, and FTY720 **(Figure 3b, Supp. Figure 6)**. Within the macrophage compartment, SA-IL-10 treatment significantly reduced the percentage of pro-inflammatory CD86^+^ F4/80^+^ CD11b^+^ macrophages (M1-like macrophages) compared to PBS, WT IL-10, and FTY720, while increasing the percentage of immunoregulatory CD206^+^ F4/80^+^ CD11b^+^ macrophages (M2-like macrophages) compared to PBS and FTY720, as well as the percentage of PD-L1^+^ F4/80^+^ CD11b^+^ macrophages compared to FTY720 **(Figure 3c-e, Supp. Figure 6)**.

**Figure 3.**
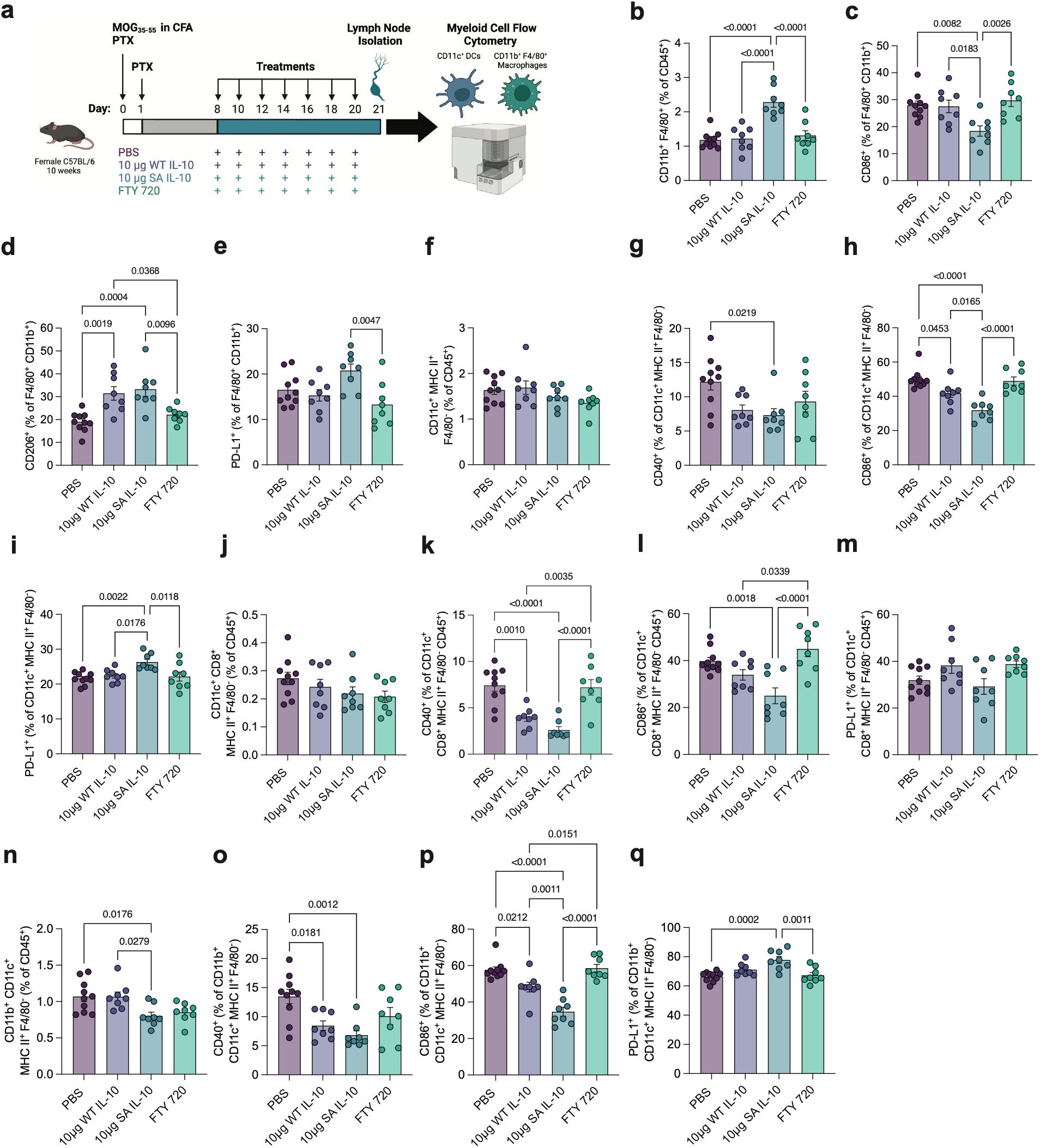
SA-IL-10 reduces the activation of pro-inflammatory myeloid cells in the spinal cord draining lymph nodes of EAE-bearing mice. **a.** Overview of experimental design. EAE-bearing mice were prophylactically administered s.c. PBS or s.c. 10 μg WT IL-10, or 10 μg SA-IL-10 (equimolar with respect to IL-10) every other day from day 8 to day 20, or i.g. FTY720 (1 mg/kg body weight) daily from day 8 as described in **Fig. 2d** (*n* = 8-10 mice/group). On day 21, the iliac and cervical LNs (spinal cord draining lymph nodes, SC-dLNs) were excised and processed for flow cytometric analysis. **b-e.** Percentage of **(b)** CD11b^+^ F4/80^+^ of CD45^+^ cells, and **(c)** CD86^+^, **(d)** CD206^+^, **(e)** PD-L1^+^ of F4/80^+^ CD11b^+^ CD45^+^ cells. f-i. Percentage of **(f)** CD11c^+^ MHC class II^+^ F4/80^-^ of CD45^+^ cells, and **(g)** CD40^+^, **(h)** CD86^+^, and (i) PD-L1^+^ of CD11c^+^ MHC class II^+^ F4/80^-^ cells. **j-m.** Percentage of (j) CD11c^+^ CD8^+^ MHC class II^+^ F4/80^-^ of CD45^+^ cells, and **(k)** CD40^+^, **(l)** CD86^+^, and **(m)** PD-L1^+^ of CD11c^+^ CD8^+^ MHC class II^+^ F4/80^-^ cells. **n-q.** Percentage of **(n)** CD11b^+^ CD11c^+^ MHC class II^+^ F4/80^-^ of CD45^+^ cells, and **(o)** CD40^+^, **(p)** CD86^+^, and **(q)** PD-L1^+^ of CD11b^+^ CD11c^+^ MHC class II^+^ F4/80^-^ cells. Data represent means ± s.e.m. Statistical analysis was performed using one-way ANOVA with Tukey’s multiple-comparisons test. Diagram in **(a)** was created using BioRender.com.

SA-IL-10 administration did not alter the percentage of CD11c^+^ MHC class II^+^ F4/80^-^ bulk DCs in the SC-dLNs **(Figure 3f, Supp. Figure 6)**. However, within the bulk DC compartment, SA-IL-10 significantly reduced the percentage of CD40^+^ DCs compared to PBS and reduced the percentage of CD86^+^ DCs compared to PBS, WT IL-10, and FTY720, while increasing the percentage of PD-L1^+^ DCs compared to PBS, WT IL-10, and FTY720 **(Figure 3g-i, Supp. Figure 6).**

SA-IL-10 did not significantly affect the percentage of CD11c^+^ CD8^+^ MHC class II^+^ F4/80^-^ cDC1s in the SC-dLNs **(Figure 3j, Supp. Figure 6)**. Within the cDC1 compartment, SA-IL-10 treatment significantly reduced the percentage of CD40^+^ cDC1s compared to PBS, WT IL-10, and FTY720 and reduced the percentage of CD86^+^ cDC1s compared to PBS and FTY720, without significantly altering the percentage of PD-L1^+^ cDC1s **(Figure 3k-m, Supp. Figure 6)**.

Given the established role of cDC2s in mediating CD4^+^ T cell priming during EAE,^47^ we next examined the impact of SA-IL-10 administration on cDC2s in the SC-dLNs. SA-IL-10 administration significantly reduced the percentage of CD11b^+^ CD11c^+^ MHC class II^+^ F4/80^-^ cDC2s compared to PBS and WT IL-10 in the SC-dLNs **(Figure 3n, Supp. Figure 6)**. Within the cDC2 compartment, SA-IL-10 significantly reduced the percentage of CD40^+^ cDC2s compared to PBS and reduced the percentage of CD86^+^ cDC2s compared to PBS, WT IL-10, and FTY720, while increasing the percentage of PD-L1^+^ cDC2s compared to PBS and FTY720 **(Figure 3o-q, Supp. Figure 6)**.

### SA-IL-10 suppresses pathogenic T_H_17 cells and increases the expression of checkpoint markers on T_H_2 cells in the SC-dLNs of EAE-bearing mice

To evaluate the impact of prophylactic SA-IL-10 treatment on T cell populations in the SLOs of EAE-bearing mice, we isolated cells from the SC-dLNs and performed immunophenotyping **(Figure 4a, Supp. Figure 7)**. SA-IL-10 administration did not significantly alter the percentage of CD4^+^ T cells in the SC-dLNs compared to PBS **(Figure 4b, Supp. Figure 7a)**. Within the CD4^+^ T cell compartment, SA-IL-10 significantly reduced the percentage of disease-associated RORγt^+^ Foxp3^-^ CD4^+^ T_H_17 cells compared to PBS and FTY720 and reduced the percentage of antigen-specific MOG tetramer^+^ RORγt^+^ Foxp3^-^ CD4^+^ T_H_17 cells compared to PBS and FTY720 **(Figure 4c-d, Supp. Figure 7a)**. SA-IL-10 did not significantly affect the frequency of Tbet^+^ Foxp3^-^ CD4^+^ T_H_1 cells, GATA-3^+^ Foxp3^-^ CD4^+^ T_H_2 cells, or Foxp3^+^ CD25^+^ CD4^+^ Tregs **(Figure 4e-g, Supp. Figure 7a)**.

**Figure 4.**
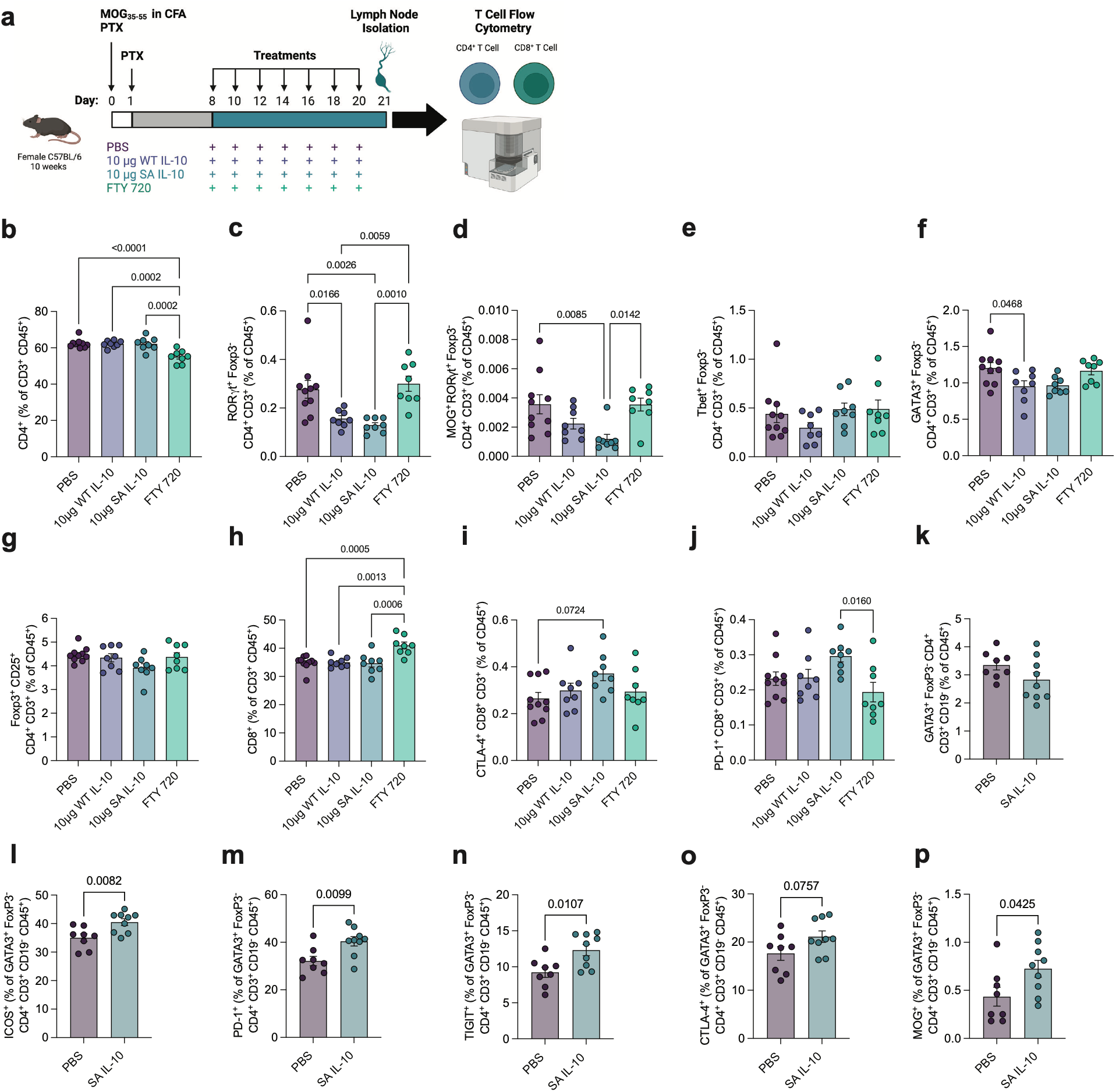
SA-IL-10 suppresses pathogenic T_H_17 cells and increases checkpoint marker expression on T_H_2 cells in the SC-dLNs of EAE-bearing mice. **a.** Overview of experimental design. EAE-bearing mice were prophylactically administered s.c. PBS or s.c. 10 μg WT IL-10, or 10 μg SA-IL-10 (equimolar with respect to IL-10) every other day from day 8 to day 20, or i.g. FTY720 (1 mg/kg body weight) daily from day 8 as described in **Fig. 2d** (*n* = 8-10 mice/group). On day 21, the SC-dLNs were excised and processed for flow cytometric analysis. **b-g.** Percentage of **(b)** CD4^+^ of CD3^+^ CD45^+^ cells, and **(c)** RORγt^+^ Foxp3^-^ CD4^+^ CD3^+^ and **(d)** MOG tetramer^+^ RORγt^+^ Foxp3^-^ CD4^+^ CD3^+^ of CD45^+^ cells, **(e)** Tbet^+^ Foxp3^-^ CD4^+^ CD3^+^, **(f)** GATA-3^+^ Foxp3^-^ CD4^+^ CD3^+^, and **(g)** Foxp3^+^ CD25^+^ CD4^+^ CD3^+^ of CD45^+^ cells. **h-j.** Percentage of **(h)** CD8^+^ of CD3^+^ CD45^+^ cells, and **(i)** CTLA-4^+^ CD8^+^ CD3^+^ and **(j)** PD-1^+^ CD8^+^ CD3^+^ of CD45^+^ cells. In an additional experiment described in **Supp. Figure 8**, the SC-dLNs were processed for flow cytometric analysis on day 19 (*n* = 8-9 mice/group). **k-o.** Percentage of **(k)** GATA-3^+^ Foxp3^-^ CD4^+^ CD3^+^ CD19^-^ of CD45^+^ cells, and **(l)** ICOS^+^, **(m)** PD-1^+^, **(n)** TIGIT^+^, **(o)** CTLA-4^+^, and **(p)** MOG tetramer^+^ of GATA-3^+^ CD4^+^ CD3^+^ CD19^-^ CD45^+^ cells. Data represent means ± s.e.m. Statistical analysis was performed using one-way ANOVA with Tukey’s multiple-comparisons test in **(b-j)** or unpaired, two-tailed Student’s t-test in **(k-p)**. Diagram in **(a)** was created using BioRender.com.

SA-IL-10 also did not significantly alter the percentage of CD8^+^ T cells in the SC-dLNs compared to PBS, WT IL-10, and FTY720 **(Figure 4h, Supp. Figure 7a)**. Within the CD8^+^ T cell compartment, SA-IL-10 treatment showed a trend toward increased frequency of CTLA-4^+^ and PD-1^+^ CD8^+^ T cells compared to PBS **(Figure 4i-j, Supp. Figure 7a)**.

In an additional experiment directly comparing PBS- and SA-IL-10 **(Supp. Figure 8a-b)**, SA-IL-10 treatment significantly increased the frequency of checkpoint markers on GATA-3^+^ Foxp3^-^ CD4^+^ T_H_2 cells in the SC-dLNs **(Figure 4k-o, Supp. Figure 7b)**. In particular, SA-IL-10 significantly increased the percentages of ICOS^+^ **(Figure 4l)**, PD-1^+^ **(Figure 4m)**, and TIGIT^+^ **(Figure 4n)** T_H_2 cells, and trended toward increasing the percentage of CTLA-4^+^ **(Figure 4o)** T_H_2 cells. SA-IL-10 also significantly increased the percentage of antigen-specific, MOG-tetramer^+^ GATA-3^+^ FoxP3^+^ T_H_2 cells **(Figure 4p)**. Upregulation of checkpoint molecules on CD4^+^ T cells has been shown to reduce proliferation and effector cytokine production.^48^ SA-IL-10 may suppress EAE in part by promoting a checkpoint-expressing T_H_2 phenotype.

### SA-IL-10 reduces antigen-induced pro-inflammatory cytokine secretion in MOG_35-55_-restimulated splenocytes from EAE-bearing mice

To characterize the impact of SA-IL-10 on antigen-induced cytokine production, we restimulated spleen-derived cells from EAE-bearing mice *ex vivo* with MOG_35-55_ peptide and measured cytokine production **(Figure 5a)**. SA-IL-10 treatment significantly suppressed IFN-γ production compared to PBS **(Figure 5b)**, IL-6 production compared to PBS and WT IL-10 **(Figure 5d)**, IL-17A production compared to PBS **(Figure 5g)**, IL-17F production compared to PBS **(Figure 5h)**, and IL-13 production compared to PBS **(Figure 5j)**, and showed a trend toward reduced IL-9 production compared to PBS **(Figure 5f)**. SA-IL-10 treatment did not significantly affect TNF-α **(Figure 5c)**, IL-4 **(Figure 5e)**, IL-22 **(Figure 5i)**, or IL-2 **(Figure 5k)** production compared to PBS or WT IL-10. Reduced IFN-γ, IL-17A, IL-17F, and IL-6 levels are consistent with the dampening of T_H_1 and T_H_17-associated cytokines.^49^ These *ex vivo* data demonstrate that SA-IL-10 treatment limits pro-inflammatory cytokine production following *ex vivo* MOG_35-55_ restimulation.

**Figure 5.**
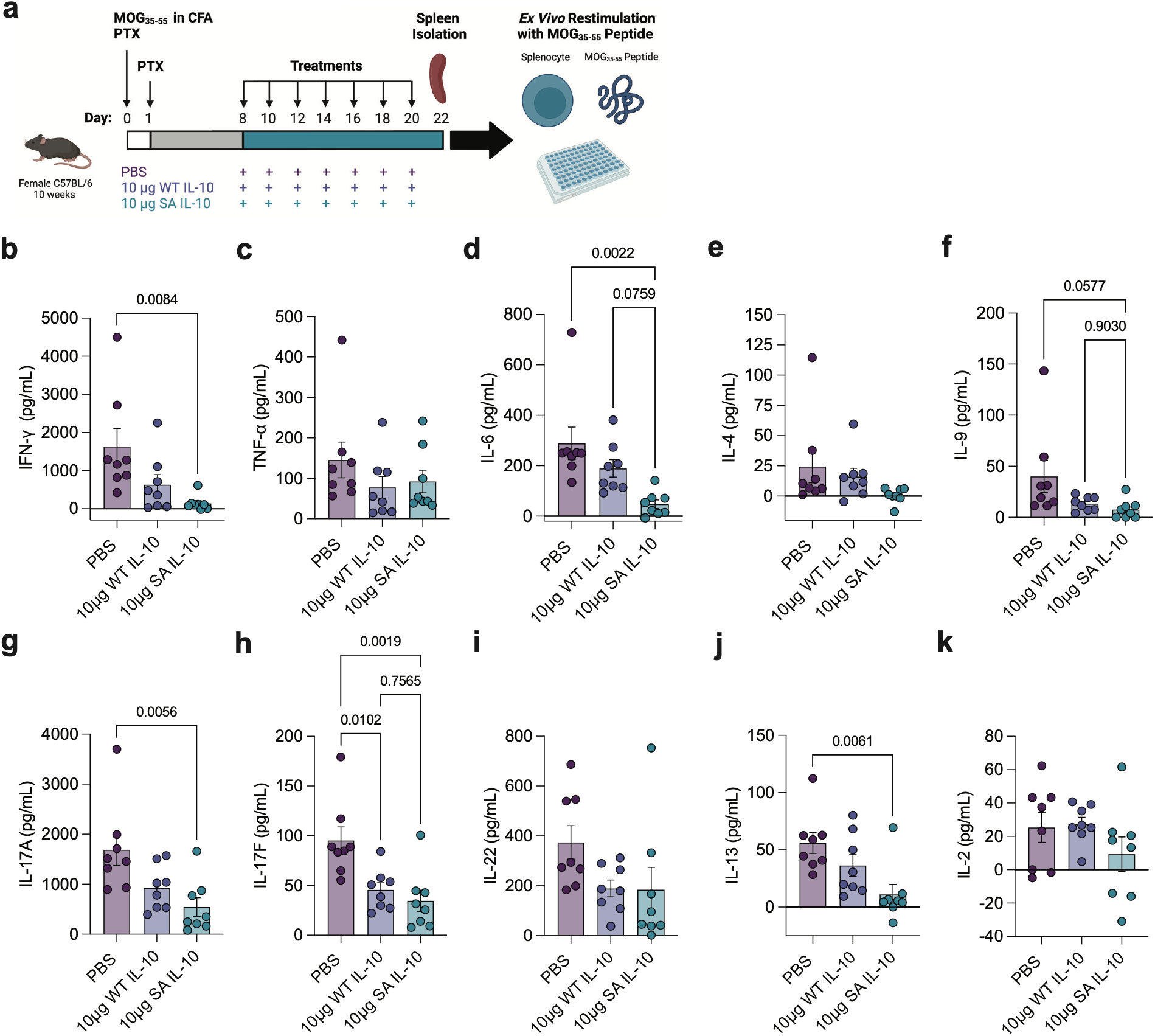
SA-IL-10 reduces antigen-induced pro-inflammatory cytokine secretion in MOG_35-55_-restimulated splenocytes from EAE-bearing mice. **a.** Overview of experimental design. EAE-bearing mice were prophylactically administered s.c. PBS, 10 μg WT IL-10, or 10 μg SA-IL-10 (equimolar with respect to IL-10) every other day from day 8 to day 20 as described in **Fig. 2a** (*n* = 8 mice/group). On day 22, the spleens from the EAE-bearing mice were excised and processed. Spleen-derived cells were restimulated *ex vivo* for 72 hr with MOG_35-55_ peptide, and cytokine secretion was quantified in the culture supernatants. **b-k.** Concentration of **(b)** IFN-γ, **(c)** TNF-α, **(d)** IL-6, **(e)** IL-4, **(f)** IL-9, **(g)** IL-17A, **(h)** IL-17F, **(i)** IL-22, **(j)** IL-13, and **(k)** IL-2 in the restimulated cell supernatant. Values represent background-subtracted cytokine levels, calculated as the difference between restimulated and unstimulated samples. Data represent means ± s.e.m. Statistical analysis was performed using one-way ANOVA with Tukey’s multiple-comparisons test. Diagram in **(a)** was created using BioRender.com.

### SA-IL-10 reduces leukocyte infiltration and activation in the spinal cords of EAE-bearing mice

A hallmark of MS and EAE is the infiltration of pathogenic leukocytes into the CNS.^44^ EAE develops when autoreactive, MOG-specific RORγt^+^ T_H_17 cells breach the disrupted blood-brain barrier and infiltrate the spinal cord, where they are reactivated, initiating an inflammatory cascade that leads to chronic hindlimb paralysis.^44^ To assess the impact of prophylactic SA-IL-10 treatment on leukocyte frequency and activation in the CNS, we isolated spinal cord-derived cells, stained them for lymphoid and myeloid markers, and performed immunophenotyping **(Figure 6a)**.

**Figure 6.**
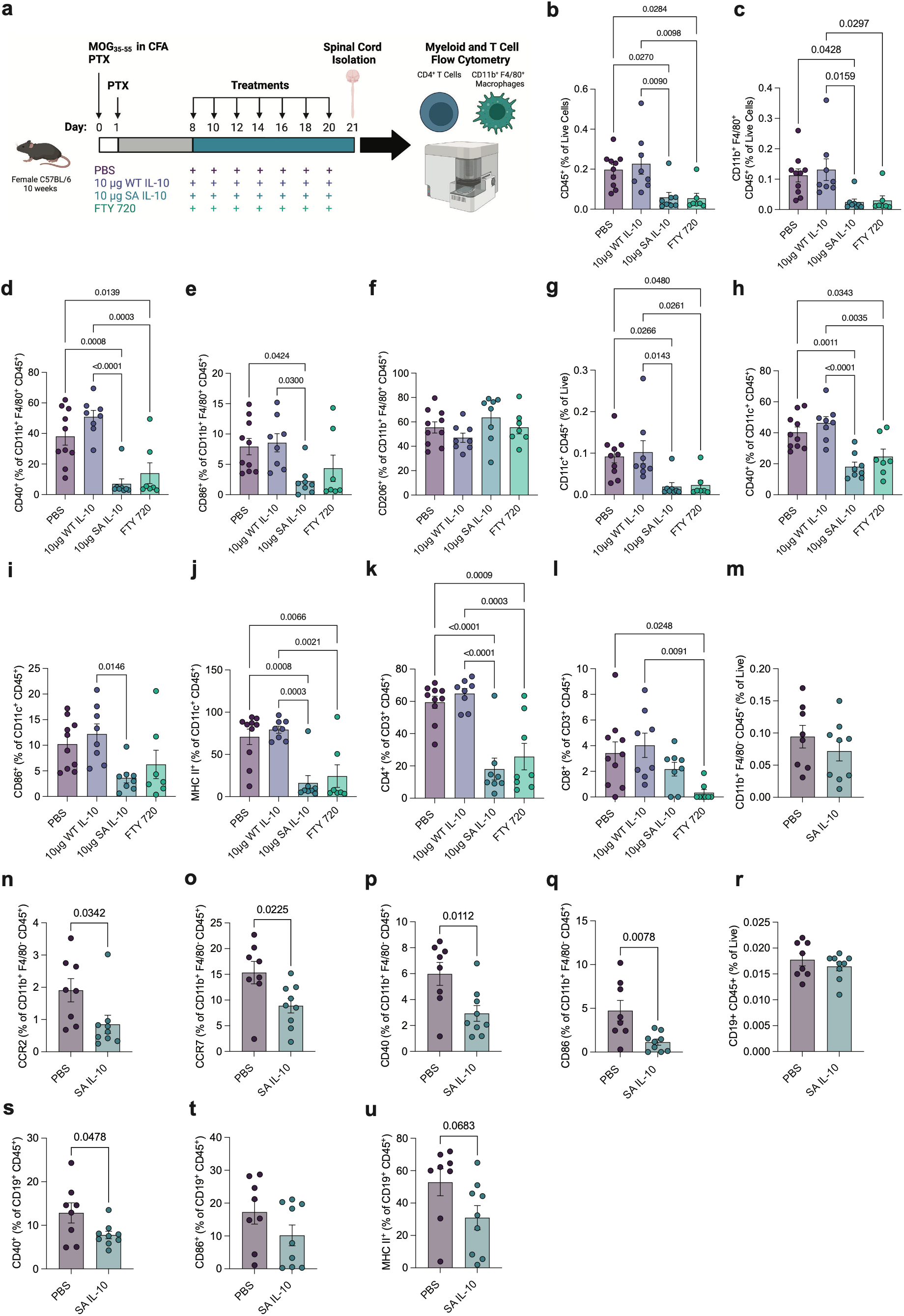
SA-IL-10 ameliorates leukocyte infiltration and activation in the spinal cords of EAE-bearing mice. **a.** Overview of experimental design. EAE-bearing mice were prophylactically administered s.c. PBS or s.c. 10 μg WT IL-10, or 10 μg SA-IL-10 (equimolar with respect to IL-10) every other day from day 8 to day 20, or i.g. FTY720 (1 mg/kg body weight) daily from day 8 as described in **Fig. 2d** (*n* = 8-10 mice/group). On day 21, the spinal cords were excised and processed for flow cytometric analysis. **b-c.** Percentage of **(b)** CD45^+^ and **(c)** CD11b^+^ F4/80^+^ CD45^+^ of live cells. **d-f.** Percentage of **(d)** CD40^+^, **(e)** CD86^+^, and **(f)** CD206^+^ of CD11b^+^ F4/80^+^ CD45^+^ cells. **g-j.** Percentage of **(g)** CD11c^+^ CD45^+^ of live cells, and **(h)** CD40^+^, **(i)** CD86^+^, and **(j)** MHC class II^+^ of CD11c^+^ CD45^+^ cells. **k-l.** Percentage of **(k)** CD4^+^ and **(l)** CD8^+^ of CD3^+^ CD45^+^ cells. In an additional experiment described in **Supp. Figure 8**, the spinal cords were processed for flow cytometric analysis on day 19 (*n* = 8-9 mice/group). **m-q.** Percentage of **(m)** CD11b^+^ F4/80^-^ CD45^+^ of live cells, and **(n)** CCR2^+^, **(o)** CCR7^+^, **(p)** CD40^+^, and **(q)** CD86^+^ of CD11b^+^ F4/80^-^ CD45^+^ cells. **r-u.** Percentage of **(r)** CD19^+^ CD45^+^ of live cells, and **(s)** CD40^+^, **(t)** CD86^+^, and **(u)** MHC class II^+^ of CD19^+^ CD45^+^ cells. Data represent means ± s.e.m. Statistical analysis was performed using one-way ANOVA with Tukey’s multiple-comparisons test in **(b-l)** or unpaired, two-tailed Student’s t-test in **(m-u)**. Diagram in **(a)** was created using BioRender.com.

In the spinal cord, SA-IL-10 treatment significantly reduced the percentage of CD45^+^ leukocytes compared to PBS and WT IL-10 **(Figure 6b)**. SA-IL-10 treatment also significantly reduced the percentage of CD11b^+^ F4/80^+^ macrophages compared to PBS and WT IL-10 **(Figure 6c)**. Within the macrophage compartment, SA-IL-10 significantly reduced the frequency of CD40 and CD86 on CD11b^+^ F4/80^+^ cells compared to PBS and WT IL-10 **(Figure 6d-e)**. SA-IL-10 treatment significantly reduced the percentage of CD11c^+^ DCs compared to PBS and WT IL-10 **(Figure 6g)**. Within the DC compartment, SA-IL-10 significantly reduced the frequency of CD40 and MHC class II on CD11c^+^ cells compared to PBS and WT IL-10 and significantly reduced the frequency of CD86 on CD11c^+^ cells compared to WT IL-10 **(Figure 6h-j)**. In the T cell compartment, SA-IL-10 treatment significantly reduced the percentage of CD4^+^ T cells in the spinal cord compared to PBS and WT IL-10 but did not significantly affect the percentage of CD8^+^ T cells **(Figure 6k-l)**.

In an additional experiment comparing PBS- and SA-IL-10-treated mice **(Supp. Figure 8)**, SA-IL-10 treatment did not significantly alter the percentage of CD11b^+^ F4/80^-^ monocytes in the spinal cord **(Figure 6m)**. However, within the monocyte compartment, SA-IL-10 significantly reduced the percentages of CCR2^+^ CD11b^+^ F4/80^-^ monocytes **(Figure 6n)** and CCR7^+^ CD11b^+^ F4/80^-^ monocytes compared to PBS **(Figure 6o)**, as well as the frequency of CD40 and CD86 on CD11b^+^ F4/80^-^ monocytes **(Figure 6p-q)**. SA-IL-10 treatment did not significantly impact the percentage of CD19^+^ B cells in the spinal cord compared to PBS **(Figure 6r)**. Within the B cell compartment, SA-IL-10 significantly reduced the frequency of CD40 on CD19^+^ cells **(Figure 6s)** and trended towards reducing the frequency of CD86 and MHC class II on CD19^+^ cells **(Figure 6t-u)**. These data indicate that SA-IL-10 treatment reduces both leukocyte infiltration as well as myeloid and B cell activation in the spinal cords of EAE-bearing mice.

### SA-IL-10-treated EAE-bearing mice exhibit greater body weight gain and plasma biochemistry parameters comparable to PBS-treated mice

To assess the tolerability of repeated SA-IL-10 administration, EAE-bearing mice were administered SA-IL-10 subcutaneously every other day beginning on day 8 after disease induction, for a total of seven injections **(Figure 7a)**. Body weight was monitored longitudinally as a clinical indicator of health. PBS- and WT IL-10-treated EAE-bearing mice exhibited transient body weight loss following disease onset, whereas SA-IL-10-treated mice gained significantly more weight over the course of the study **(Figure 7b)**, consistent with reduced disease burden and suggesting that repeated SA-IL-10 administration was well tolerated.

**Figure 7.**
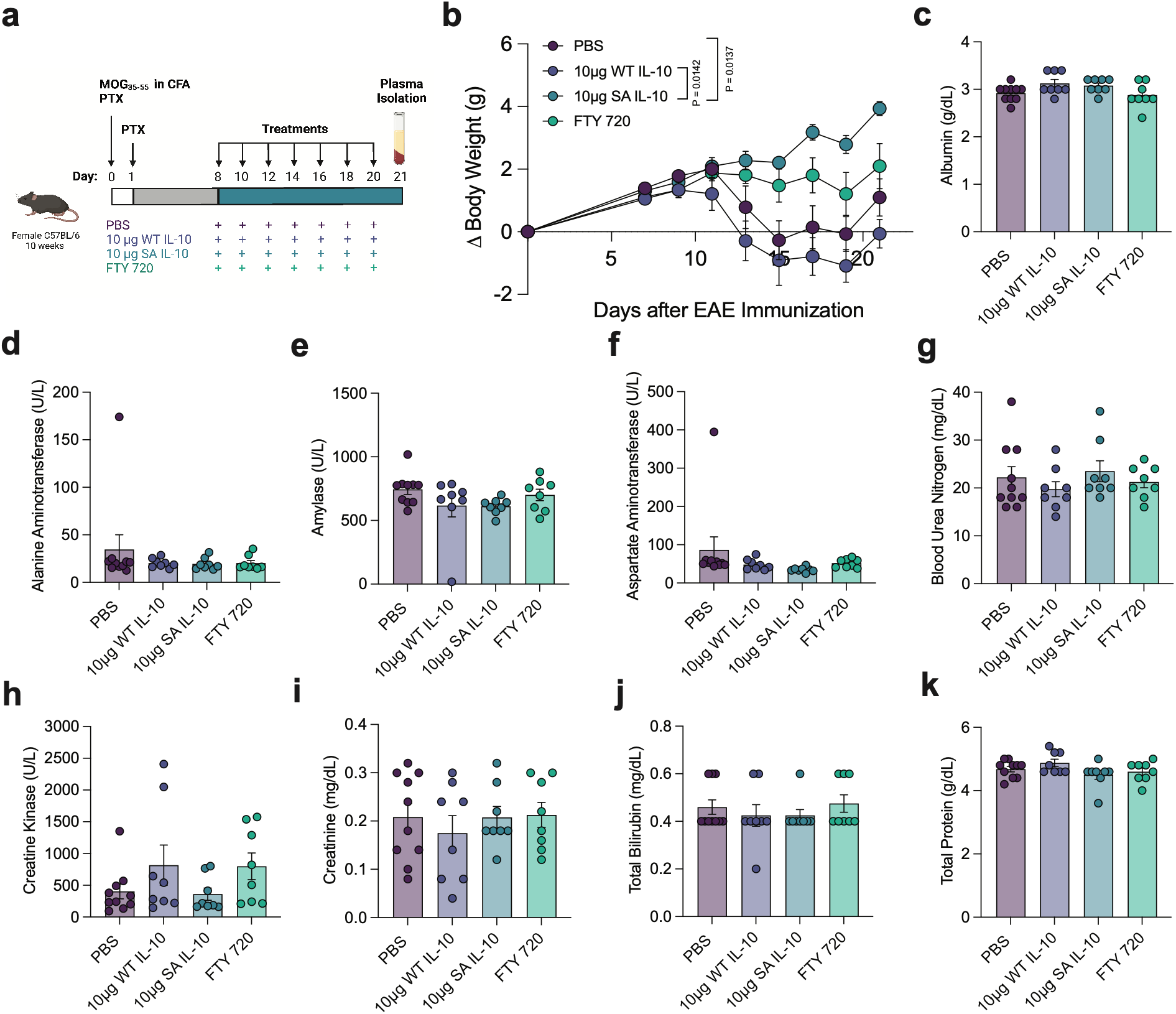
SA-IL-10-treated EAE-bearing mice exhibit greater body weight gain and plasma biochemistry parameters comparable to PBS-treated controls. **a.** Overview of experimental design. EAE-bearing mice were prophylactically administered s.c. PBS, s.c. 10 μg WT IL-10, or 10 μg SA-IL-10 (equimolar with respect to IL-10) every other day from day 8 to day 20, or i.g. FTY720 (1 mg/kg body weight) daily from day 8 as described in **Fig. 2d** (*n* = 8-10 mice/group). On day 21, blood was collected and processed for plasma biochemistry analysis. **b.** Body weight change over the course of treatment. **c-k.** Plasma concentration of **(c)** albumin, **(d)** alanine aminotransferase, **(e)** amylase, **(f)** aspartate aminotransferase, **(g)** blood urea nitrogen, **(h)** creatine kinase, **(i)** creatinine, **(j)** total bilirubin, and **(k)** total protein. Data represent means ± s.e.m. Statistical analysis was performed using one-way ANOVA with Tukey’s multiple-comparisons test. Diagram in **(a)** was created using BioRender.com.

To evaluate potential systemic toxicity associated with repeated SA-IL-10 dosing, blood was collected on day 21 after disease induction and analyzed for plasma biochemistry parameters **(Figure 7a)**. SA-IL-10 administration did not significantly alter the levels of albumin, alanine aminotransferase, amylase, aspartate aminotransferase, blood urea nitrogen, creatine kinase, creatinine, total bilirubin, and total protein compared to PBS, WT IL-10, or FTY720 **(Figure 7c-k)**. These markers serve as biochemical indicators of hepatic, renal, cardiac, and pancreatic function^50^. The absence of differences across treatment groups suggests that the administered dose and dosing regimen of SA-IL-10 did not induce detectable organ toxicity in EAE-bearing mice at this time point. These findings are consistent with our previous study demonstrating that a single dose of SA-IL-10 did not induce organ toxicity in healthy mice.^11^

## Discussion

In this study, we evaluated the effects of sustained IL-10 exposure in the SLOs using EAE as a murine model of neuroinflammation. We show that subcutaneous administration of SA-IL-10, a long-circulating fusion of SA and the immunoregulatory cytokine IL-10, prevented the development and progression of MOG_35-55_-induced EAE. In EAE, SA-IL-10 demonstrated superior prophylactic efficacy to WT IL-10 and comparable protection to FTY720, an FDA-approved MS therapy. In the SLOs, SA-IL-10 administration suppressed pathogenic T_H_17 cells, reduced myeloid cell activation, and dampened antigen-induced, pro-inflammatory cytokine production, while expanding immunoregulatory myeloid populations and increasing the percentage of checkpoint-expressing T_H_2 cells. SA-IL-10 administration also reduced leukocyte infiltration and activation in the spinal cord. These findings demonstrate that enhancing IL-10 exposure in the SLOs via albumin fusion promotes immunoregulation and prevents the onset of neuroinflammatory disease in this murine model.

Our scRNA-seq analysis identified a higher percentage of *IL10RA*-expressing Treg CD4^+^ T cells in the CSF of treatment-naïve MS patients relative to IIH controls. In contrast, *IL10* was detected in a lower percentage of Treg CD4^+^ T cells as well as non-Treg CD4^+^ T cells and DCs in both MS patients and IIH controls. These findings suggest that CSF CD4^+^ T cells in MS patients may retain the capacity to respond to IL-10, while endogenous *IL10* expression in these populations is limited. Because these analyses are based on transcriptomic data, they do not directly assess IL-10 protein levels or downstream signaling activity. Our interpretation is also limited by the patient cohort size (*n* = 5 in each of the MS and IIH control cohorts).^41^ Larger patient cohorts would be required to further investigate the extent of IL-10 axis dysregulation in MS. Incorporating protein-level measurements in future studies would also be important to determine how IL-10 cytokine and receptor expression contribute to downstream signaling in MS.

WT IL-10 has shown variable efficacy in preclinical models of neuroinflammation, with outcomes ranging from suppression of disease to no significant effect on disease, depending on the species, strain, and disease induction method.^36,37,39^ These different results are attributed, in part, to the short half-life and limited tissue persistence of WT IL-10 following systemic administration.^3,10,11^ We previously demonstrated that recombinant fusion of SA to IL-10 prolongs its plasma half-life and enhances its accumulation in the SLOs compared to WT IL-10, because of the extended circulation conferred by albumin fusion.^11^ These enhanced pharmacokinetic properties provided a rationale for testing SA-IL-10 in EAE to examine how sustained IL-10 exposure in the SLOs shapes immune responses during neuroinflammation. Prophylactic SA-IL-10 administration prevented the development and progression of MOG_35-55_-induced EAE with superior efficacy to WT IL-10. This improvement in disease prevention motivated us to next evaluate the impact of SA-IL-10 on immune cell populations within the SLOs and spinal cord.

As the SC-dLNs are key sites of antigen presentation and T cell priming, myeloid cells regulate pathogenic T_H_1 and T_H_17 differentiation in EAE.^49^ Depletion of macrophages has been shown to attenuate clinical disease severity, implicating myeloid cells in EAE pathogenesis.^51^ In the SC-dLNs, SA-IL-10 treatment reduced the frequency of CD86^+^ M1-like macrophages and increased the frequency of CD206^+^ M2-like macrophages among CD11b^+^ F4/80^+^ cells. In bulk DCs as well as in cDC1 and cDC2 subsets, SA-IL-10 treatment reduced the frequency of the co-stimulatory molecules CD40 and CD86, while increasing the frequency of the tolerogenic marker PD-L1 on bulk DCs and cDC2s. These changes in myeloid activation marker expression suggest that SA-IL-10 treatment may reduce the co-stimulatory capacity of antigen-presenting cells to prime pathogenic effector T cells in the SC-dLNs. This interpretation aligns with previous studies reporting that IL-10 reduces macrophage and DC activation marker expression.^4,52,53^

In addition to suppressing pro-inflammatory myeloid cells, SA-IL-10 treatment also modulated T cell phenotype in the SC-dLNs. IL-17A-producing CD4^+^ T cells have been shown to express IL-10Rα *in vivo*, suggesting that T_H_17 cells may respond to IL-10 signaling.^54^ In the SC-dLNs of SA-IL-10-treated EAE-bearing mice, we observed a significant reduction in the frequency of RORγt^+^ T_H_17 cells, including antigen-specific MOG tetramer^+^ RORγt^+^ T_H_17 cells, compared to PBS- and FTY720-treated controls. This reduction was supported by a significant decrease in T_H_17-associated cytokine production (IL-17A, IL-17F, and IL-6) by spleen-derived cells from SA-IL-10-treated mice following *ex vivo* restimulation with MOG_35-55_ peptide. Although the frequency of Tbet^+^ T_H_1 T cells in the SC-dLNs was not altered by SA-IL-10 administration, IFN-γ production by spleen-derived cells from SA-IL-10-treated mice was significantly reduced following *ex vivo* restimulation, suggesting that SA-IL-10 may also dampen T_H_1 effector function. Consistent with these findings, previous studies have shown that blockade of IL-10 signaling expands T_H_17 cells during intestinal inflammation^54^ and that IL-10-deficient mice exhibit a systemic increase in T_H_17 cells compared to WT mice.^55^ Additionally, myelin-specific T cells stimulated under T_H_17-polarizing conditions have also been shown to produce IL-10, which suppressed their pathogenicity and attenuated EAE in adoptive transfer models.^56,57^ These findings indicate that SA-IL-10 reduces pathogenic T cell responses, particularly T_H_17 cells, in the SLOs of EAE-bearing mice.

We also observed significant increases in the percentages of ICOS-, PD-1-, and TIGIT-expressing GATA-3^+^ T_H_2 cells and a trend towards increase in the percentage of CTLA-4-expressing T_H_2 cells in the SC-dLNs of SA-IL-10-treated EAE-bearing mice. Because SA-IL-10 did not impact the overall frequency of GATA-3^+^ T_H_2 cells, these findings suggest that SA-IL-10 treatment altered the phenotype of the T_H_2 compartment. ICOS contributes to T_H_2 differentiation and function by increasing IL-4 production and IL-4 receptor signaling.^58–60^ Higher ICOS surface expression has also been associated with increased IL-10 production,^58,61^ whereas ICOS deficiency increases susceptibility to EAE.^60^ In contrast, PD-1, TIGIT, and CTLA-4 can limit T cell effector responses by inhibiting TCR-mediated activation, proliferation, and cytokine production.^62–65^ Disruption of these inhibitory pathways has been reported to exacerbate EAE and increase immune cell infiltration into the CNS.^62,63,66,67^ Although PD-1, TIGIT, and CTLA-4 are highly expressed on Foxp3^+^ Tregs, Foxp3^-^ IL-10-producing CD4^+^ T cell populations that co-express these inhibitory receptors have also been described.^68^ The elevated frequencies of ICOS-, PD-1-, and TIGIT-expressing GATA-3^+^ T_H_2 T cells following SA-IL-10 treatment suggest that SA-IL-10 may promote an immunoregulatory T_H_2 phenotype that could contribute to limiting pathogenic inflammation in EAE. Future studies will investigate whether this population has immunosuppressive functions by characterizing its cytokine-production profile and ability to suppress responder T cell activation in *ex vivo* co-culture assays.

Pathogenic leukocyte infiltration into the CNS is a hallmark of MS and EAE.^44^ Immunophenotyping of the spinal cord revealed that SA-IL-10 treatment reduced the percentages of CD45^+^ leukocytes, including CD11b^+^ F4/80^+^ macrophages, CD11c^+^ DCs, and CD4^+^ T cells. SA-IL-10 treatment also reduced the frequency of the activation markers CD40, CD86, and MHC class II on CNS-infiltrating myeloid cells. During EAE, myeloid cells upregulate chemokine receptors that facilitate migration into the CNS.^14^ SA-IL-10 treatment reduced the frequency of CCR7^+^ and CCR2^+^ monocytes in the spinal cord, suggesting diminished recruitment and/or retention of inflammatory monocytes in the CNS. These findings indicate that prophylactic SA-IL-10 administration may limit immune cell infiltration and activation in the CNS.

Prophylactic SA-IL-10 treatment demonstrated comparable efficacy to FTY720, a clinically approved MS therapy. FTY720 acts as a functional antagonist of sphingosine 1-phosphate receptors, causing receptor internalization and thereby restricting lymphocyte egress from the SLOs.^46^ Although broad immunosuppression can limit pathogenic immune responses, it may also interfere with endogenous immunoregulatory pathways.^14^ In addition to reducing pathogenic RORγt^+^ T_H_17 cells, SA-IL-10 treatment also increased the frequency of CD206^+^ M2-like macrophages, indicating a shift toward an immunoregulatory myeloid phenotype. FTY720 has been reported to increase IL-10 production by human monocyte-derived DCs and B cells in MS patients, suggesting that IL-10-associated mechanisms may also contribute to its therapeutic effects.^21,31,32^ Given the overlapping yet distinct immunomodulatory effects of SA-IL-10 and FTY720, future studies could investigate whether combined treatment provides greater protection than either therapy alone.

A key limitation of cytokine-based therapies is the toxicity caused by the high and frequent dosing required to compensate for short circulation half-life.^12^ Unmodified recombinant human IL-10 has shown limited clinical efficacy and dose-dependent adverse effects, including flu-like symptoms, hematologic abnormalities, and thrombocytopenia, in multiple clinical studies.^10,69^ These outcomes suggest that the rapid clearance, systemic exposure, and insufficient tissue persistence of WT IL-10 limit its therapeutic window. In contrast, in the present study, SA-IL-10 was well tolerated in EAE-bearing mice at the dose and dosing frequency administered. SA-IL-10-treated mice gained significantly more body weight over the course of treatment and exhibited no significant differences in plasma biochemistry markers indicative of hepatic, renal, and pancreatic function compared to PBS-treated controls. Due to its improved tissue retention, SA-IL-10 may reduce the need for frequent or high-dose administration, potentially mitigating toxicity observed with WT IL-10 in prior clinical trials. Together with the immunomodulatory effects observed here, these safety findings support the continued preclinical evaluation of SA-IL-10.

In conclusion, this study demonstrates that sustained IL-10 exposure in the SLOs modulates immune responses during neuroinflammation. Using EAE as a murine model, we show that subcutaneous administration of SA-IL-10 suppresses pathogenic T cell and myeloid populations in the SLOs, and reduces immune cell infiltration and activation in the spinal cord. SA-IL-10 treatment also induced the upregulation of multiple checkpoint molecules on T_H_2 T cells. Human scRNA-seq analysis identified higher percentages of *IL10RA-*expressing Treg CD4^+^ T cells in the CSF of treatment-naïve MS patients compared to IIH controls, alongside limited *IL10* gene expression in these cells, suggesting that this population may retain the capacity to respond to IL-10 while expressing low levels of the cytokine gene. These data motivate supplementation with IL-10 engineered for prolonged circulation. Given its extended half-life, immunomodulatory effects in the SLOs and CNS, and favorable safety profile, SA-IL-10 may be a promising therapeutic strategy for neuroinflammatory diseases.

### Limitations of this study

In this study, we focused on prophylactic administration of SA-IL-10 to prevent the onset of MOG_35-55_-induced EAE. SA-IL-10 was evaluated primarily for its ability to prevent disease development in this model. Because most MS patients initiate treatment after symptom onset, future studies will need to evaluate the effects of therapeutic SA-IL-10 administration during the chronic phase of disease. Additionally, as MS occurs in both chronic and relapsing-remitting forms, it will be important to evaluate SA-IL-10 in relapsing-remitting models, such as the PLP_131-151_ model, to determine its effects on relapse frequency and disease severity.

## Materials and Methods

### Study design

The goal of this study was to genetically fuse SA to IL-10 to improve its persistence in the SLOs and evaluate its prophylactic efficacy in EAE, a murine model of neuroinflammation. Recombinant SA-IL-10 was expressed and evaluated *in vivo* alongside WT IL-10 and FTY720. EAE disease development and progression were monitored longitudinally. Immune cell phenotypes in the SLOs and spinal cord were assessed via flow cytometry, and antigen-specific cytokine responses were evaluated by *ex vivo* restimulation assays. The tolerability of SA-IL-10 treatment was studied by monitoring body weight and plasma biochemistry parameters. Sample size was determined based on previous and preliminary studies.^43,70^ Mice were randomly assigned to treatment groups. Disease scoring was performed by an investigator blinded to the treatment groups and prior scores.

### Single-cell RNA sequencing data analysis

A previously published single-cell transcriptomic dataset was obtained from the NCBI Gene Expression Omnibus (accession GSE138266).^41,71^ The dataset initially comprised PBMC and CSF immune cells from treatment-naïve MS patients (*n* = 6) and IIH control patients (*n* = 6).^41^ One MS CSF sample (pseudonym 58637) and one IIH control CSF sample (pseudonym 45044) were excluded from the analysis due to lack of published paired PBMC data.^41,71^ The resulting analyses were performed on paired CSF and blood cells from five treatment-naïve MS patients (*n* = 5) and five IIH control patients (*n* = 5). All downstream analyses were performed using *R* (v. 4.5.1) with *Seurat* (v. 5.3.0).

Quality control was performed based on the number of detected genes per cell (*nFeature*), total unique molecular identifier (UMI) counts per cell (*nCount*), and the percentage of transcripts mapping to mitochondrial genes (percent.mt). Cells were retained if they contained 200 to 5,000 detected genes, 500 to 25,000 UMIs, and no more than 15% mitochondrial gene transcripts. We used *DropletUtils* to process one sample which was deposited as a raw matrix (CSF for PST83775). Genes detected in fewer than three cells were removed from the expression matrix. Doublets were identified and removed using *scDblFinder* (v. 1.22.0) with an expected doublet rate of 7.5%, yielding 61,228 cells (38,980 PBMCs, 22,248 CSF cells). The dataset was then split by tissue type (PBMC and CSF). For downstream analyses, PBMC and CSF samples were processed and analyzed independently. The two compartments were not integrated.

Each tissue object was normalized using *SCTransform* (v. 0.4.2), utilizing variable regression of mitochondrial transcripts (*percent.mt*) and selection of 3,000 variable features. Principal component analysis was performed using 50 components. Samples were then integrated across patients within each tissue using *Harmony* (v. 1.2.4) with theta = 2 and a maximum of 20 iterations. A shared nearest-neighbor graph was constructed using the first 30 Harmony dimensions (*FindNeighbors*, dim = 30), and clusters were identified (*FindClusters*) using the Smart Local Moving algorithm at a resolution of 1.5. UMAP plots were computed on the first 30 Harmony dimensions for visualization.

Cell-type annotation was performed at the cluster-level. Cluster identities were first assigned by reference-based annotation run on log-normalized counts with *SingleR* (v. 2.10.0), using the Monaco Immune Reference Dataset (*celldex* v. 1.18.0)^42^ with broad-grained labels. Per-cluster label confidence was assessed from the *SingleR* score heatmap, and clusters with annotation confidence scores below 0.6 were designated “Unlabeled.” Clusters identified as T cells in broad-grained labeling (Monaco labels = CD4^+^ T cells, T cells) were further resolved using the Monaco Immune Reference Dataset with fine-grained reference labels. Fine-grained reference labels of the T cells were grouped into the following subsets for downstream analysis: Non-Treg CD4^+^ T cells, Treg CD4^+^ T cells, CD8^+^ T cells, and *γδ* T cells. Final cluster labels were verified against dot plot visualization of canonical lineage markers: Non-Treg CD4^+^ T cells (*CD3D*, *CD3E*, *TRAC, CD4*), Treg CD4^+^ T cells (*CD3D*, *CD3E, TRAC, CD4, FOXP3*, *IL2RA*, *CTLA4*, *IKZF2*, *TIGIT*), CD8^+^ T cells (*CD8A*, *CD8B*, *CD3D*, *CD3E*, *TRAC*), *γδ* T cells (*TRDC, TRGC1, TRGC2, CD3D, CD3E,*), B cells (*MS4A1, CD19, IGHD*), natural killer (NK) cells (*NKG7, PRF1, GNLY, GZMB*), monocytes (*CD14, MS4A7, S100A9, LYZ*), dendritic cells (*HLA-DRA*, *FLT3, CLEC4A, ITGAX*), and progenitors (*CD34, KIT, SPINK2*).

Downstream analyses of the IL-10 pathway focused on monocytes, DCs, non-Treg CD4^+^ T cells, and Tregs. Expression of *IL10*, *IL10RA*, and *IL10RB* genes was visualized as UMAP feature plots of SCT-normalized values and as violin plots of log-normalized per-cell expression values. For quantitative patient-level analyses, a cell was considered positive for a gene if at least one UMI mapped to that gene. The percentage of cells expressing each gene was calculated separately for each patient, cell population, and tissue.

### Production and purification of recombinant proteins

To generate WT IL-10 and SA-IL-10, codon optimized DNA sequences encoding mouse IL-10 or mouse SA without its pro-peptide (amino acids 25-608), mouse IL-10, and a (GGGS)_2_ linker **(Supp. Table 2)** were synthesized and subcloned into the mammalian expression vector pcDNA3.1(+) by GenScript, as previously described.^11,43^ A C-terminal hexahistidine (His_6_) tag was included to enable affinity purification. In both constructs, a single cysteine residue within the murine IL-10 was substituted with tyrosine to reduce disulfide-mediated dimerization during expression and purification, as described previously.^72^ The recombinant proteins were expressed in suspension-adapted HEK 293-F cells (Gibco) cultured in FreeStyle 293 Medium (Gibco), as previously described.^11,43^ On the day of transfection, HEK 293-F cells were resuspended in fresh FreeStyle 293 medium at a density of 1 x 10^6^ cells mL^-1^. A mixture of 1 μg mL^-1^ plasmid DNA and 2 μg mL^-1^ linear 25 kDa polyethyleneimine (Polysciences) prepared in OptiPRO Serum-Free Medium (Gibco) was sequentially added to suspension cells. Transfected cells were then incubated at 37°C in 5% CO_2_ with orbital shaking at 135 rpm. After 7 days of culture, the supernatant was collected, centrifuged, passed through a 0.22 μm filter (Corning), and stored at 4°C until purification.

Affinity and size exclusion chromatography (SEC) purification was carried out using an ÄKTA Pure 25M (Cytiva), as previously described.^11,43^ Briefly, affinity chromatography was performed by loading the supernatant into a 5 mL HisTrap HP column (Cytiva). The column was then washed with binding buffer (20 mM NaH_2_PO_4_, 0.5 M NaCl, 20 mM imidazole, all from Sigma Aldrich, at pH 7.4). His_6_-tagged proteins were removed from the column by flowing a gradient of elution buffer (500 mM imidazole, 20 mM NaH_2_PO_4_, 0.5 M NaCl at pH 7.4). The proteins were then loaded into a HiLoad Superdex 200 Prep Grade column (Cytiva), further purified by SEC, and eluted in phosphate buffered saline (PBS, 137 mM NaCl, 2.7 mM KCl, 10 mM Na_2_HPO_4_, and 1.8 mM KH_2_PO_4_). All purification steps were performed at 4°C. The purified proteins were confirmed to contain < 0.01 endotoxin units mL^-1^ via a HEK-Blue TLR-4 reporter assay. The concentration of the recombinant proteins was determined by measuring their absorbance at 280 nm using a NanoDrop spectrophotometer (Thermo Scientific).

### Mice

Female, 8-week-old C57BL/6 mice were purchased from Charles River Laboratories. Mice were acclimated at the University of Chicago Animal Facility for 2 weeks prior to use. Animals were housed on a 12-hour light/dark cycle at room temperature (20–24°C). All experiments and procedures were performed in accordance with the University of Chicago Institutional Animal Care and Use Committee.

### MOG_35-55_ EAE model

EAE was induced as previously described^43,70^ by subcutaneously immunizing female, 10-week-old, C57BL/6 mice (Charles River) with MOG_35-55_ antigen emulsified in Complete Freund’s Adjuvant in the dorsal flank, followed by intraperitoneal administration of pertussis toxin from *Bordetella pertussis* (Hooke Laboratories) dissolved in PBS on days 0 and 1, according to manufacturer’s instructions. Disease score was determined daily from day 7 according to a scoring criterion established by Hooke Laboratories. Mice were administered subcutaneous PBS, WT IL-10, or SA-IL-10 in the dorsal flank every other day, or intragastric (i.g.) FTY720 (1 mg/kg body weight) daily from day 8 until the experimental endpoint. Treatment groups were randomly assigned across cages. Disease scores were monitored daily, and body weight was recorded every other day beginning on day 8 by an investigator blinded to treatment groups and prior measurements.

### Preparation of single-cell suspensions from EAE tissues

At the experimental endpoint, the spinal cord, cervical and iliac lymph nodes (spinal cord draining lymph nodes, SC-dLNs), and spleen were harvested. These tissues were incubated with Collagenase D (2 mg mL^-1^, Roche) in DMEM (Gibco) supplemented with 10% heat-inactivated fetal bovine serum (FBS, Gibco) and 1.2 mM CaCl_2_ (Sigma-Aldrich) for 45 minutes at 37°C, as previously described.^11,43^ The tissues were then mechanically dissociated and passed through a sterile, 70 μm filter (Corning). Red blood cells in spleen suspensions were lysed using ACK Lysing Buffer (Gibco) for 5 minutes and quenched with 25 mL DMEM supplemented with 10% FBS, as previously described.^11,43^ Single-cell suspensions were then washed once with DMEM and resuspended in complete RPMI 1640 medium, consisting of RPMI 1640 medium supplemented with 10% FBS, 1% Penicillin-Streptomycin (Gibco), and 50 μM β-mercaptoethanol (Sigma-Aldrich) for downstream flow cytometry staining and *ex vivo* restimulation assays.

### Flow cytometry staining of EAE tissues

Single-cell suspensions from the SC-dLNs and spinal cords were seeded in 96-well plates, washed once with PBS, and incubated with live/dead dye and Fc block (BioLegend) in PBS at 4°C for 20 minutes. Cells were washed once with FACS buffer, consisting of PBS, 2% FBS, 2 mM EDTA (Invitrogen), and stained with surface antibodies **(Supp. Table 3)** in FACS buffer for 45 minutes at 4°C. To detect MOG-reactive cells, cells were washed once in PBS and stained with APC-conjugated MOG_38-49_ Tetramer (NIH Tetramer Core Facility) in PBS for 45 minutes at 37°C. Cells were then fixed and permeabilized for 1 hour at 4°C (FoxP3/Transcription Factor Kit, eBioscience), washed twice, and stained overnight at 4°C with intracellular antibodies **(Supp. Table 3)**. Following staining, cells were washed twice and resuspended in FACS buffer. Data were acquired on a NovoCyte Penteon (Agilent) or an Aurora (Cytek) and analyzed in FlowJo (v. 10.8.0, BD Biosciences).

### Ex vivo restimulation of spleen-derived cells from EAE-bearing mice with MOG_35-55_ peptide

Spleen-derived cells (5.0 x 10^5^ cells per well) were resuspended in complete RPMI 1640 medium and seeded into 96-well plates. Cells were restimulated with 10 μM mouse MOG_35–55_ peptide (Genscript) for 72 hours at 37°C with 5% CO_2_, as previously described^43,70^. After incubation, cells were centrifuged at 1,000 x g for 5 minutes. Cell supernatants were collected for cytokine analysis using a LEGENDplex assay (BioLegend).

### Isolation of plasma from EAE-bearing mice

On day 21 after EAE induction, blood was collected from EAE-bearing mice treated with PBS, WT IL-10, SA-IL-10, or FTY720 for plasma biochemistry analysis. Samples were collected into EDTA-coated tubes (BD Biosciences) and centrifuged at 1,000 x g for 10 minutes to isolate the plasma, which was then transferred to protein LoBind tubes (Eppendorf).

### Biochemistry analysis of plasma from EAE-bearing mice

Plasma samples from mice treated with PBS, WT IL-10, SA-IL-10, or FTY720 were analyzed for albumin, alanine aminotransferase, amylase, aspartate aminotransferase, blood urea nitrogen, creatine kinase, creatinine, total bilirubin, and total protein using a biochemistry analyzer (Alfa Wassermann Diagnostic Technologies), following the manufacturer’s instructions.

### Statistical analysis

Human scRNA-seq analyses were performed using *R* (v. 4.5.1), and murine statistical analyses were performed using Prism software (v. 9, GraphPad). Unless otherwise indicated, murine data represent mean ± s.e.m. Sample sizes and statistical tests are displayed in the figure legends. In human single-cell RNA-seq analyses, the percentage of cells expressing each gene was compared between MS patients and IIH controls at the patient-level using two-sided Wilcoxon rank-sum tests with Holm-Bonferroni correction for multiple comparisons within each gene and tissue. Adjusted *P* values are displayed on the graphs for comparisons with adjusted *P* < 0.05. Hedges’ *g*, a measure of effect size suited to small clinical cohorts, was calculated using the *effsize* package (v. 0.8.1)^73,74^ and is reported with scaled 95% confidence intervals in **Supp. Table 1**. Sample sizes for murine studies were determined based on previous and preliminary studies.^43,70^ For comparisons among three or more murine experimental groups, one-way ANOVA followed by Tukey’s multiple-comparisons test was used, unless otherwise specified. For comparison of two murine experimental groups, an unpaired, two-tailed Student’s *t*-test was used, unless otherwise specified. During preparation of code for visualization of scRNA-seq data, the authors utilized Claude Fable 5 (Anthropic), Claude Opus 5 (Anthropic), and Codex 5.5 (OpenAI) to assist in writing code in the R language. After using these models, the authors reviewed and edited the code and take full responsibility for the content of the publication.

### Study approval

All animal studies were performed in accordance with protocols approved by the Institutional Animal Care and Use Committee at the University of Chicago and complied with all relevant institutional and national guidelines for the care and use of laboratory animals. For human studies, this work utilized previously published, de-identified scRNA-seq data (GSE138266).^41,71^ No new human subjects were recruited for this study, and no identifiable human data or personal identifiers were used.

## Supporting information

Budina et al Supplementary Materials 20260911

## Resource Availability

Further information and requests for data, resources, and reagents should be directed to and will be fulfilled by the corresponding author, Jeffrey A. Hubbell. Availability of the engineered cytokine is subject to production capacity and the absence of an external centralized repository. The material may be shared upon reasonable request, subject to availability and cost recovery for production, processing, and shipping.

## Code Availability

All code used to analyze scRNA-seq data in this study is publicly available at https://github.com/jwreda/il10-in-treatment-naive-ms. Additional analyses were conducted using established commercial and open-source software packages, as detailed in the Materials and Methods section.

## Author Contributions

J.A.H. oversaw all research. E.B., K.C.R, and J.A.H. designed the experiments. E.B., K.C.R., B.T.B., and L.R.V. prepared materials. E.B., J.W.R., K.C.R., B.T.B, H.C., T.N.B, A.S., M.N., S.R., C.R.F., I.V., A.L.L., K.H., S.G., and J.I. performed experiments. E.B., J.W.R., and J.L. analyzed the data. E.B., J.W.R., J.L., and J.A.H. wrote the manuscript. All authors contributed to and approved the manuscript.

## Acknowledgements

We thank Dr. A. Reder, Dr. E. Yuba, Dr. A. Mansurov, and Dr. A. Esser-Kahn for helpful discussions and review of our manuscript. We thank the University of Chicago Cytometry and Antibody Technology Core Facility, the University of Chicago Human Tissue Resource Center, and the University of Chicago Integrated Light Microscopy Core Facility (Cancer Center Support Grant P30CA014599). We thank the NIH Tetramer Core Facility (NIH Contract 75N93020D00005 and RRID:SCR_026557) for providing reagents (I-Ab, mouse MOG_38-49_, GWYRSPFSRVVH, APC).

