## Supplementary material for "Albumin-fused interleukin-10 prevents experimental autoimmune encephalomyelitis by promoting immunoregulation in the secondary lymphoid organs": Budina et al Supplementary Materials 20260911

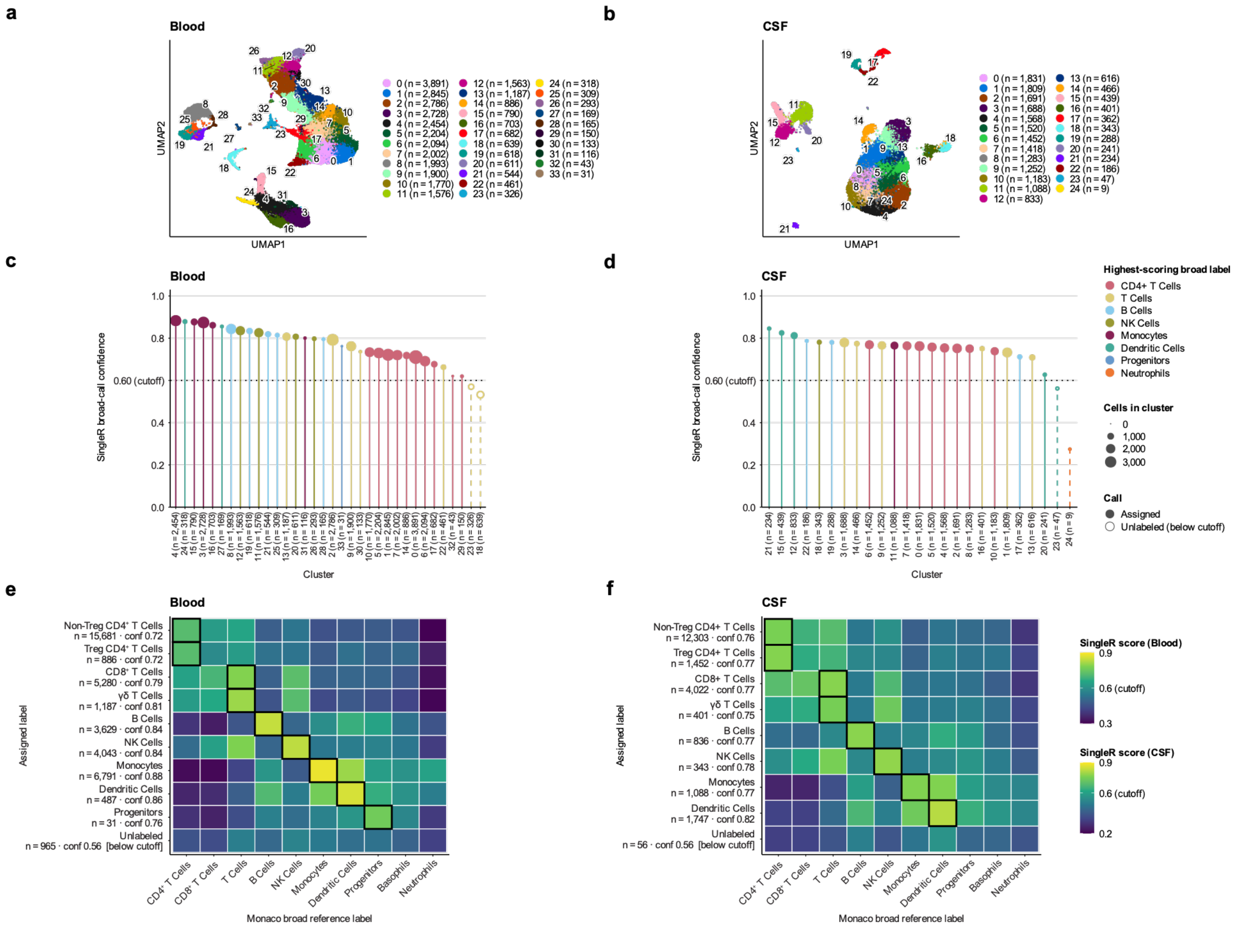


**Supplementary Figure 1 | Clustering and annotation confidence of cluster-level annotations of immune populations in blood and CSF of MS and IIH control patients.** Related to **Figure 1**. **a-b.** UMAP projections of clusters identified in **(a)** blood and **(b)** cerebrospinal fluid (CSF) annotated with number of cells per cluster. **c-d.** Lollipop plots depicting absolute *SingleR* confidence scores assigning Monaco broad labels for each cluster in blood **(c)** and CSF **(d)**. Circle size reflects cluster membership size while color reflects assigned label. The dotted line demonstrates the confidence cutoff of 0.6. **e-f.** Heatmaps of absolute *SingleR*'s confidence scores assigning a given Monaco broad label across clusters labeled in **(e)** blood and **(f)** CSF with independent scales per tissue. Rows reflect labels as displayed in **Fig. 1a-b** and columns reflect broad labels from the Monaco reference. Rows are annotated with number of cells with that label (n) and confidence of annotation ("conf"). Black outlines designate labels that are both assigned and have confidence scores above the cutoff of 0.6. Clusters that scored below this threshold for all broad labels were designated "Unlabeled." Single-cell RNA sequencing data were obtained from dataset GSE138266 (MS patients, $n$ = 5; IIH control patients, $n$ = 5).[41] Cell-type annotation was performed at the cluster level against the Monaco Immune Reference Dataset.[42]

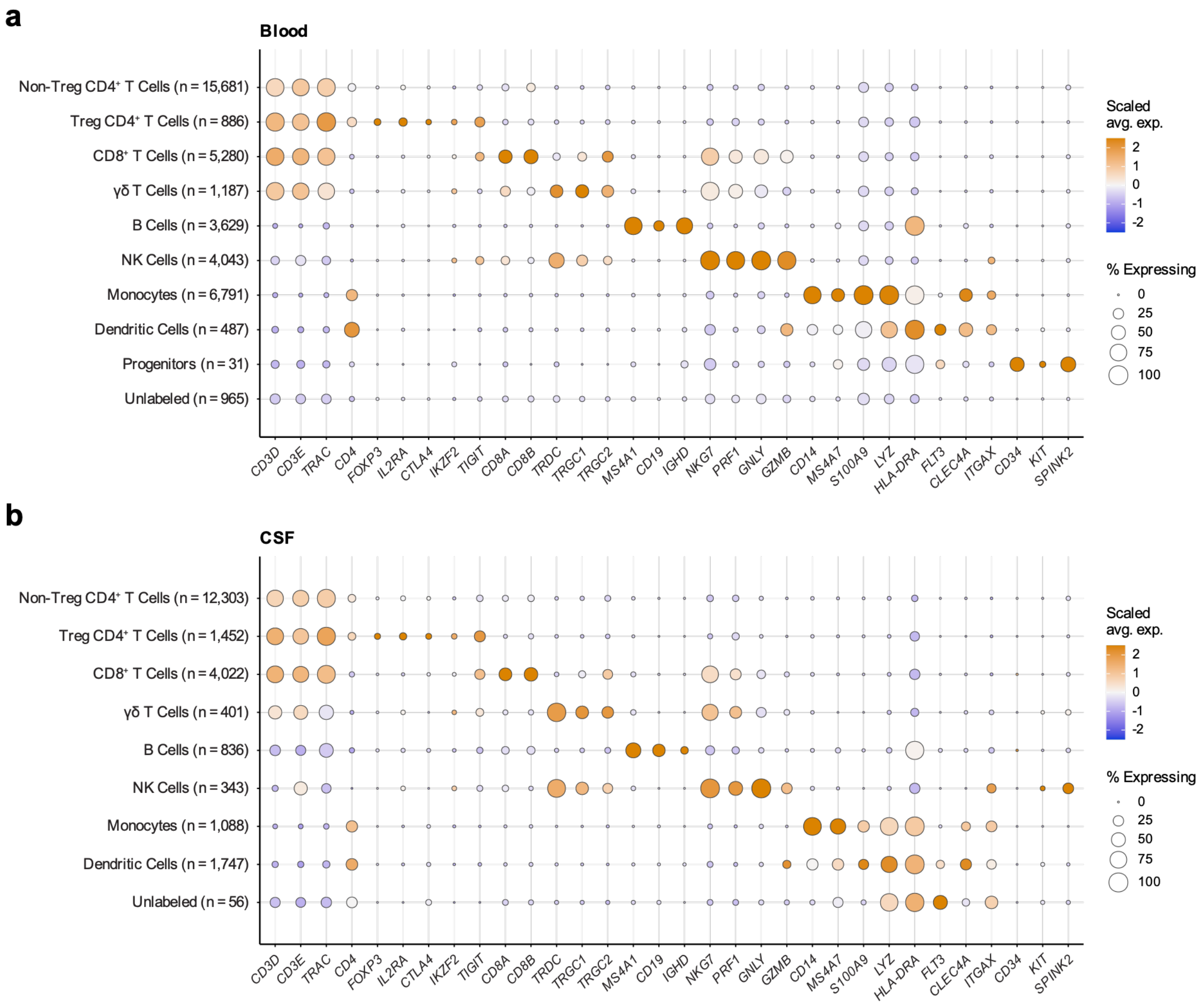


**Supplementary Figure 2 | *SingleR* annotations identify immune populations that demonstrate expression of canonical markers.** Related to **Figure 1**. **a-b.** Dot plots of canonical lineage marker genes across immune cell populations in **(a)** blood and **(b)** cerebrospinal fluid (CSF). Dot size indicates the percentage of cells within each population expressing a given gene. Dot color indicates the mean expression scaled across the immune populations (z-score, clipped at $\pm$ 2.5). Marker genes were selected *a priori* from established immune lineage definitions and grouped by lineage: *CD3D, CD3E, TRAC, CD4* (non-Treg CD4$^+$ T cells); *FOXP3, IL2RA, CTLA4, IKZF2, TIGIT* (Treg CD4$^+$ T cells); *CD8A, CD8B* (CD8$^+$ T cells); *TRDC, TRGC1, TRGC2,* ($\gamma\delta$ T cells); *MS4A1, CD19, IGHD* (B cells); *NKG7, PRF1, GNLY, GZMB* (NK cells); *CD14, MS4A7, S100A9, LYZ* (monocytes); *HLA-DRA, FLT3, CLEC4A, ITGAX* (dendritic cells); and *CD34, KIT,* and *SPINK2* (progenitors). Single-cell RNA sequencing data were obtained from dataset GSE138266 (MS patients, *n* = 5; IIH control patients, *n* = 5).[41] Cell-type annotation was performed at the cluster level against the Monaco Immune Reference Dataset,[42] with a confidence threshold of 0.6; clusters that scored below this threshold for all Monaco broad labels were designated "Unlabeled."

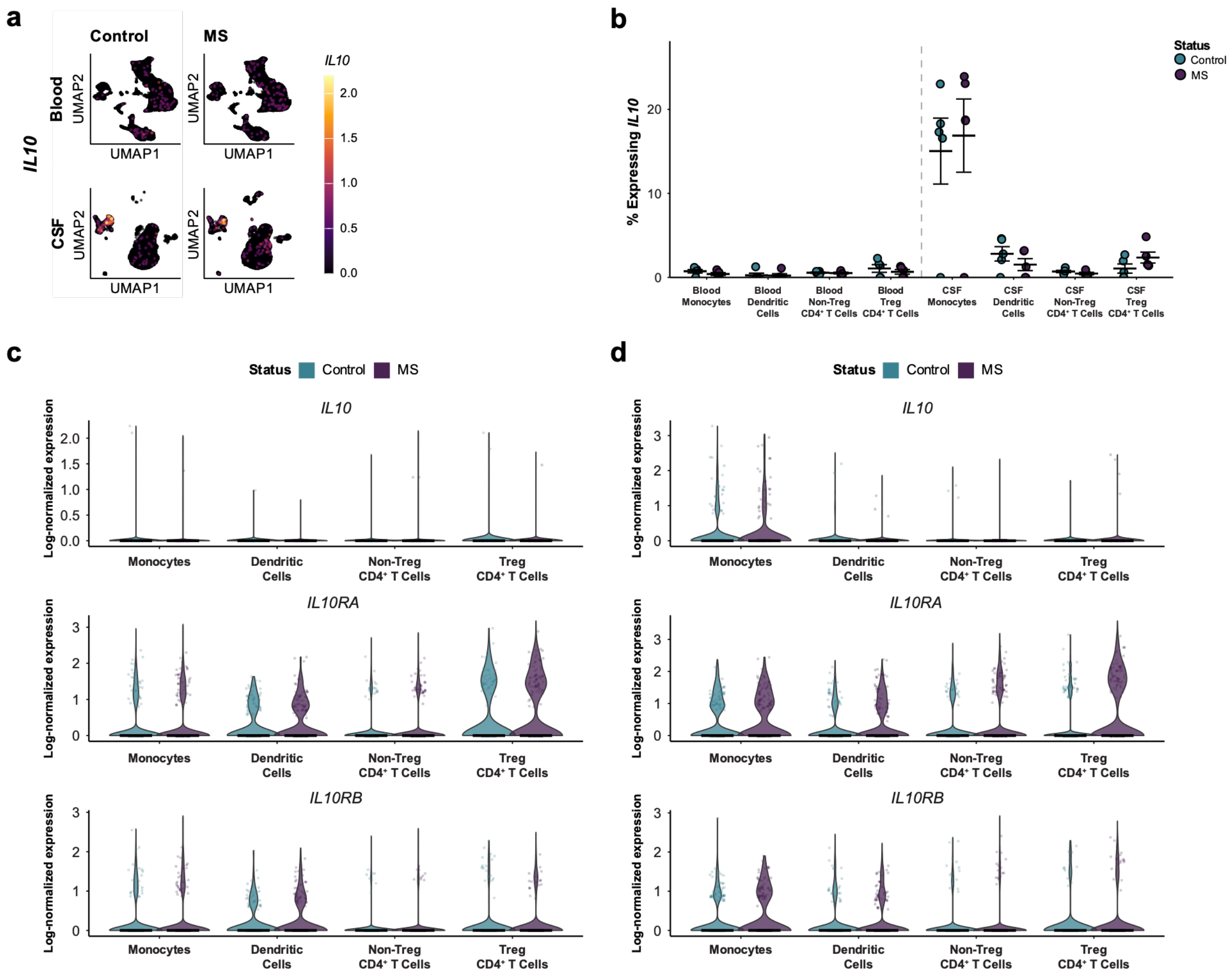


**Supplementary Figure 3 | *IL10*, *IL10RA*, and *IL10RB* gene expression across immune populations in the blood and CSF of MS and IIH control patients.** Related to **Figure 1c-f**. **a.** UMAP feature plot of SCT-normalized *IL10* gene expression levels in blood- and cerebrospinal fluid (CSF)-derived immune cells from multiple sclerosis (MS) patients and control patients with idiopathic intracranial hypertension (IIH). **b.** Percentage of monocytes, dendritic cells, non-Treg CD4$^+$ T cells, and Treg CD4$^+$ T cells expressing the *IL10* gene in the blood and CSF. **c-d.** Violin plots of log-normalized *IL10*, *IL10RA*, and *IL10RB* gene expression in monocytes, dendritic cells, non-Treg CD4$^+$ T cells, and Treg CD4$^+$ T cells from patients with multiple sclerosis (MS; purple) and control patients with idiopathic intracranial hypertension (Control; teal), shown separately for **(c)** blood and **(d)** CSF. Each violin shows the distribution of per-cell log-normalized expression values across cells pooled from all patients within the indicated diagnosis, tissue, and cell population. Overlaid points represent up to 150 individual cells, and the horizontal bar indicates the median of the pooled per-cell values. As cells are nested within patients, these plots are presented for descriptive visualization and were not used for statistical inference. Patient-level comparisons for *IL10RA* and *IL10RB* genes are reported in **Figure 1d** and **Figure 1f**. Single-cell RNA sequencing data were obtained from dataset GSE138266 (MS patients, $n$ = 5; IIH control patients, $n$ = 5).[41] Cell-type annotation was performed at the cluster level against the Monaco Immune Reference Dataset,[42] with a confidence threshold of 0.6; clusters that scored below this threshold for all Monaco broad labels were designated "Unlabeled."

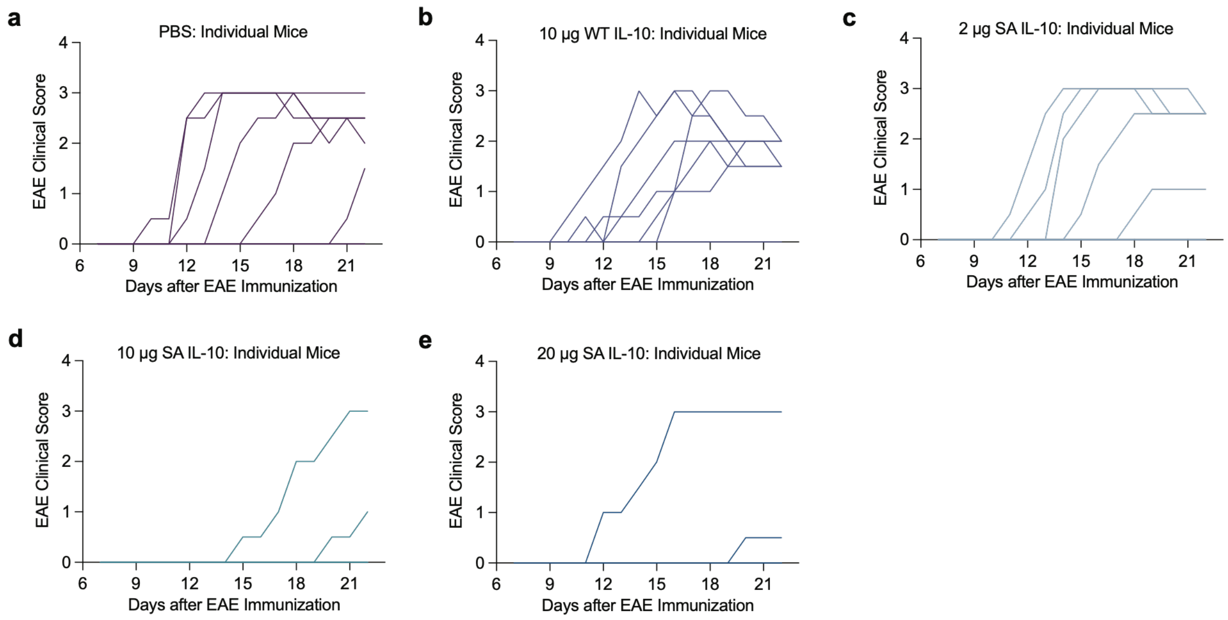


**Supplementary Figure 4 | Individual clinical scores of EAE-bearing mice in SA-IL-10 dosing study.** Related to **Figure 2a-c**. EAE-bearing mice were administered s.c. **(a)** PBS, **(b)** 10 µg WT IL-10, **(c)** 2 µg SA-IL-10, **(d)** 10 µg SA-IL-10, or **(e)** 20 µg SA-IL-10 (all equimolar with respect to IL-10) every other day from day 8 to day 20, as described in **Figure 2a** (*n* = 8 mice/group). Clinical scores were recorded daily from day 7 through the study endpoint.

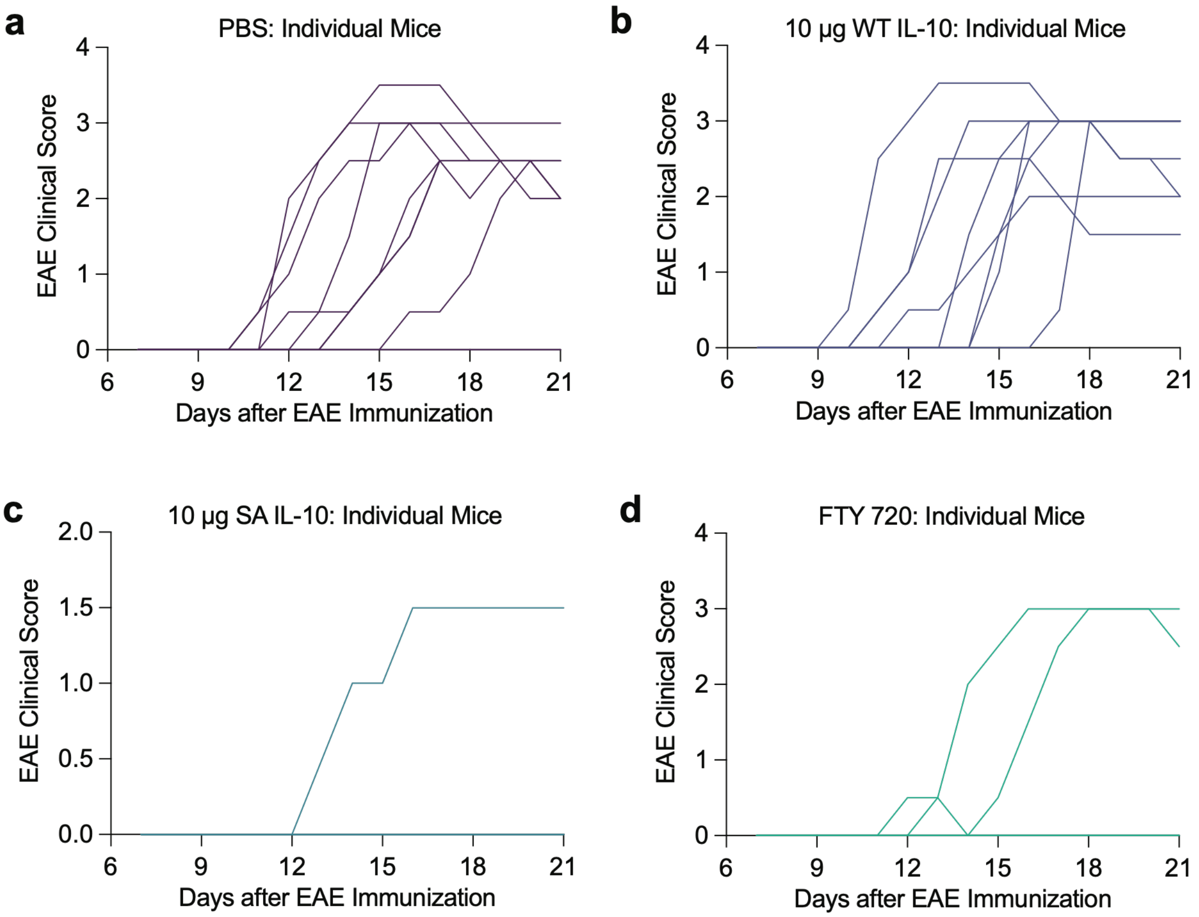


**Supplementary Figure 5 | Individual clinical scores of EAE-bearing mice in the SA-IL-10 versus FTY720 comparison study.** Related to **Figure 2d-f**. EAE-bearing mice were administered s.c. **(a)** PBS, **(b)** 10 µg WT IL-10, or **(c)** 10 µg SA-IL-10 (all equimolar with respect to IL-10) every other day from day 8 to day 20, or i.g. **(d)** FTY720 (1 mg/kg) daily from day 8 to day 20 (*n* = 8-10 mice/group), as described in **Figure 2d**. Clinical scores were recorded daily from day 7 through the study endpoint.

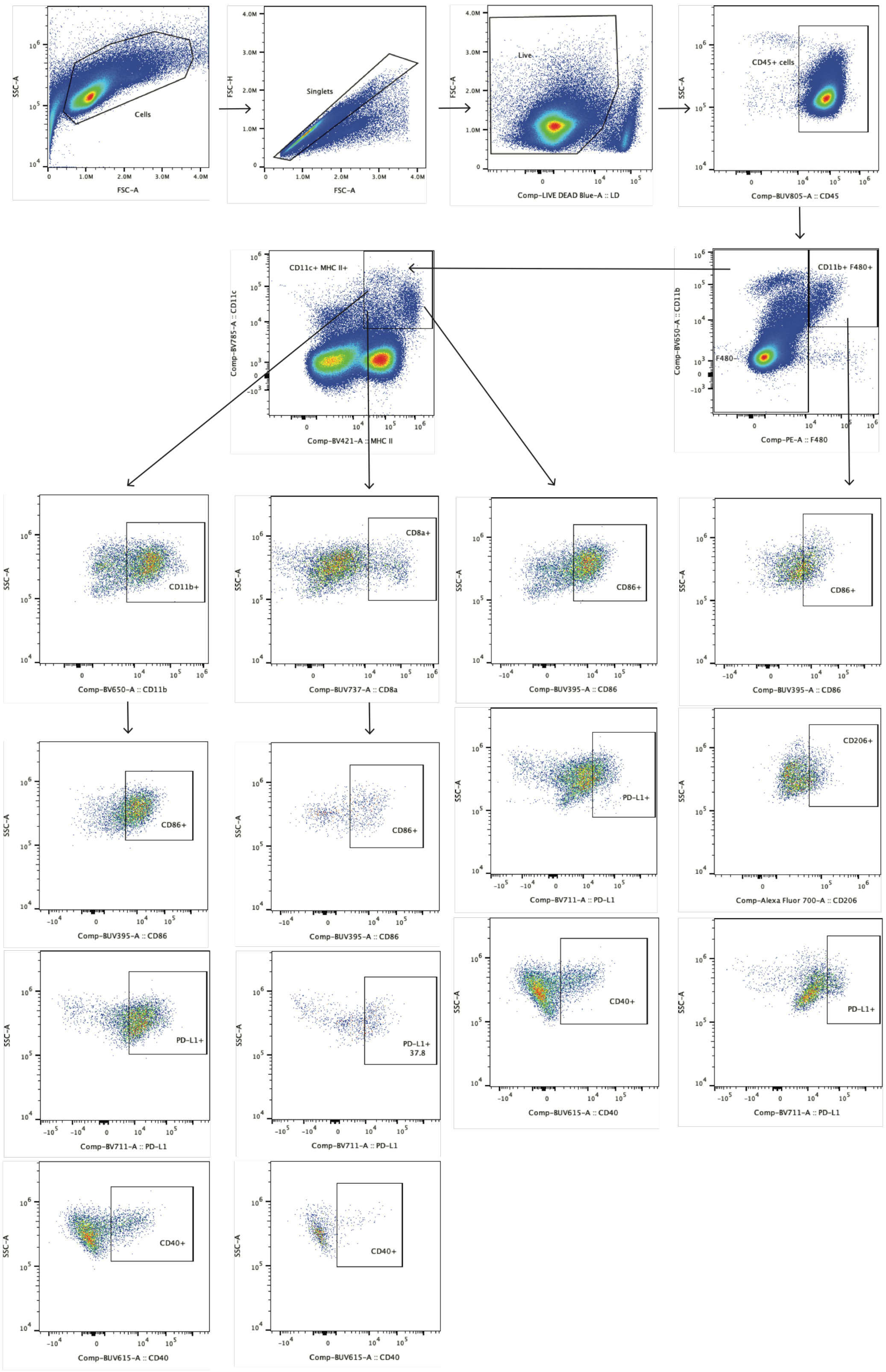


**Supplementary Figure 6 | Flow cytometry gating strategy for identification of myeloid cell populations.** Related to **Figures 3b-q** and **6b-j**. A representative gating strategy from the SC-dLNs of EAE-bearing mice is shown.

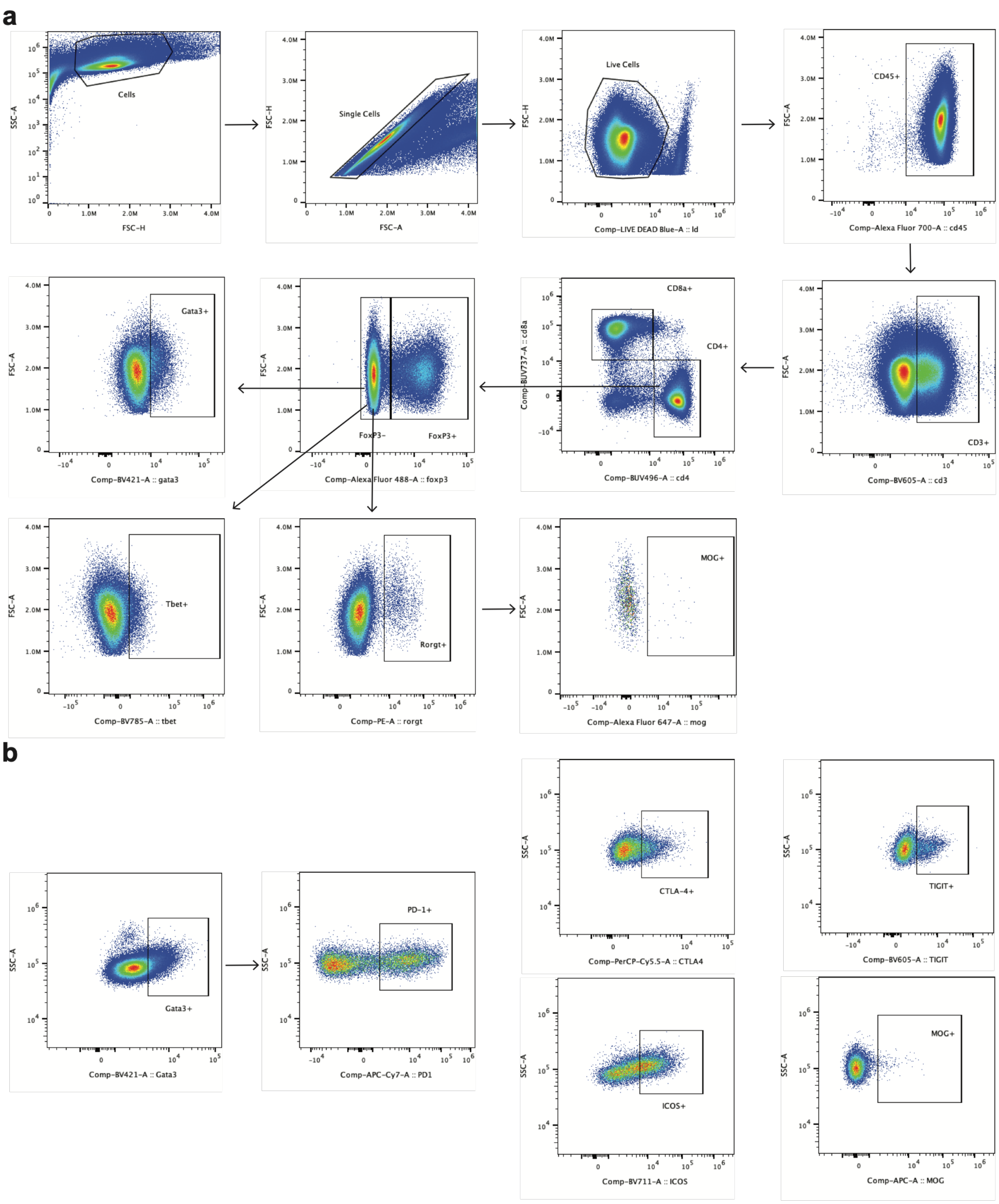


**Supplementary Figure 7 | Flow cytometry gating strategy for identification of T cell populations.** Related to **Figures 4b-o** and **6k-l**. A representative gating strategy from the SC-dLNs of EAE-bearing mice is shown. The gating strategy depicts **(a)** $CD4^+$ and $CD8^+$ T cell subsets and **(b)** checkpoint marker-expressing GATA-$3^+$ $Foxp3^-$ $CD4^+$ T cells.

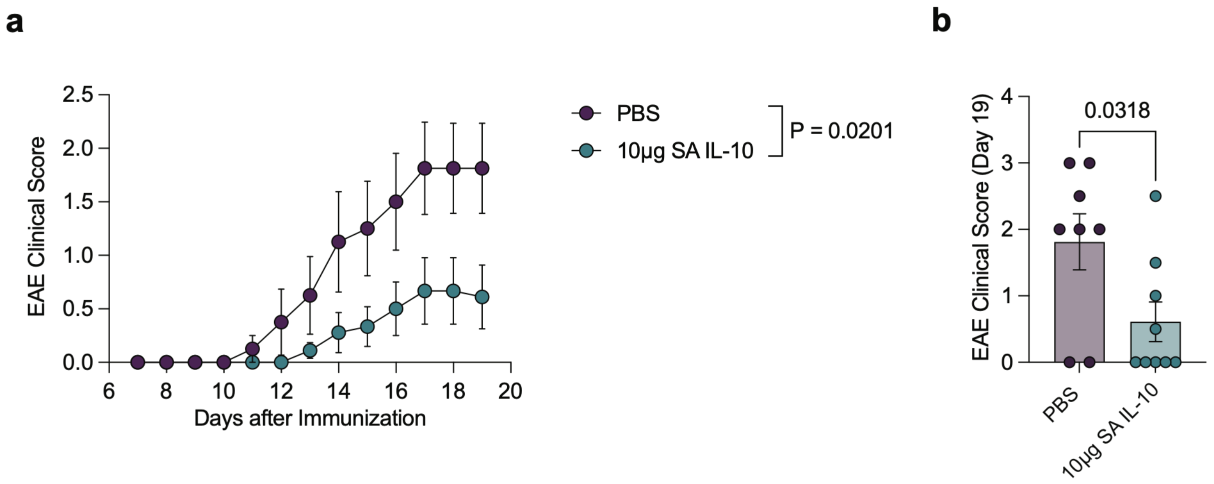


**Supplementary Figure 8 | Prophylactic SA-IL-10 administration prevents the development and progression of EAE in a repeated experiment.** Related to **Figure 2a-f**. EAE was induced in C57BL/6 mice by s.c. immunization with $MOG_{35-55}$/CFA followed by i.p. injections of PTX on days 0 and 1. Mice were prophylactically administered s.c. PBS or 10 µg SA IL-10 (equimolar with respect to IL-10) every other day from day 8 to day 18 (*n* = 8-9 mice/group). Clinical scores were recorded daily from day 7 through the study endpoint. **a.** Disease progression. **b.** Disease scores at endpoint (day 19). Data represent means ± s.e.m. Statistical analysis of disease score area-under-the-curve (from d8 to 19 in **(a)**) and EAE clinical score on day 19 in **(b)** was performed using an unpaired, two-tailed Student's t-test.

| Gene | Tissue | Cell Type | n (MS) | n (Ctrl) | Mean % MS | Mean % Ctrl | Wilcoxon padj | Hedges' g [95% CI] |
|---|---|---|---|---|---|---|---|---|
| *IL10* | Blood | Monocytes | 5 | 5 | 0.4 | 0.7 | 0.997 | -0.73 [-1.88, 0.46] |
| *IL10* | Blood | Dendritic Cells | 5 | 5 | 0.2 | 0.3 | 1.000 | -0.06 [-1.18, 1.06] |
| *IL10* | Blood | Non-Treg CD4+ T cells | 5 | 5 | 0.5 | 0.6 | 1.000 | -0.21 [-1.33, 0.92] |
| *IL10* | Blood | Tregs | 5 | 5 | 0.7 | 1.1 | 1.000 | -0.43 [-1.55, 0.72] |
| *IL10* | CSF | Monocytes | 5 | 5 | 16.9 | 15.0 | 0.889 | 0.18 [-0.95, 1.30] |
| *IL10* | CSF | Dendritic Cells | 5 | 5 | 1.5 | 2.8 | 0.889 | -0.66 [-1.81, 0.52] |
| *IL10* | CSF | Non-Treg CD4+ T cells | 5 | 5 | 0.5 | 0.7 | 0.889 | -0.71 [-1.86, 0.48] |
| *IL10* | CSF | Tregs | 5 | 5 | 2.4 | 1.1 | 0.889 | **0.88** [-0.34, 2.05] |
| *IL10RA* | Blood | Monocytes | 5 | 5 | 24.9 | 25.0 | 1.000 | -0.01 [-1.13, 1.11] |
| *IL10RA* | Blood | Dendritic Cells | 5 | 5 | 39.2 | 26.0 | 0.444 | **0.88** [-0.34, 2.06] |
| *IL10RA* | Blood | Non-Treg CD4+ T cells | 5 | 5 | 18.3 | 10.3 | 0.222 | **1.51** [0.14, 2.81] |
| *IL10RA* | Blood | Tregs | 5 | 5 | 42.3 | 30.9 | 0.286 | **1.18** [-0.11, 2.40] |
| *IL10RA* | CSF | Monocytes | 5 | 5 | 37.5 | 31.1 | 0.548 | 0.32 [-0.82, 1.44] |
| *IL10RA* | CSF | Dendritic Cells | 5 | 5 | 36.5 | 23.2 | 0.187 | **1.29** [-0.02, 2.54] |
| *IL10RA* | CSF | Non-Treg CD4+ T cells | 5 | 5 | 30.4 | 20.7 | 0.167 | **1.73** [0.30, 3.09] |
| *IL10RA* | CSF | Tregs | 5 | 5 | 43.6 | 20.6 | ***0.032*** | **2.79** [1.01, 4.51] |
| *IL10RB* | Blood | Monocytes | 5 | 5 | 19.9 | 22.2 | 0.841 | -0.32 [-1.44, 0.82] |
| *IL10RB* | Blood | Dendritic Cells | 5 | 5 | 28.2 | 14.1 | 0.286 | **1.22** [-0.07, 2.46] |
| *IL10RB* | Blood | Non-Treg CD4+ T cells | 5 | 5 | 8.3 | 4.8 | 0.127 | **1.56** [0.18, 2.88] |
| *IL10RB* | Blood | Tregs | 5 | 5 | 14.9 | 8.0 | 0.286 | **1.10** [-0.16, 2.31] |
| *IL10RB* | CSF | Monocytes | 5 | 5 | 28.5 | 20.4 | 0.927 | 0.45 [-0.70, 1.58] |
| *IL10RB* | CSF | Dendritic Cells | 5 | 5 | 25.0 | 18.8 | 0.667 | **0.94** [-0.29, 2.13] |
| *IL10RB* | CSF | Non-Treg CD4+ T cells | 5 | 5 | 9.8 | 10.7 | 0.927 | -0.29 [-1.41, 0.85] |
| *IL10RB* | CSF | Tregs | 5 | 5 | 16.1 | 8.6 | 0.381 | **1.29** [-0.02, 2.54] |

**Supplementary Table 1 | Statistical analysis of IL-10 pathway gene expression across immune cell populations.** Related to **Figure 1c-f** and **Supplementary Figure 3a-b**. Summary of statistical comparisons of the percentage of cells in monocytes, dendritic cells, non-Treg CD4+ T cells, and Treg CD4$^{+}$ T cells expressing *IL10*, *IL10RA*, and *IL10RB* genes between patients with multiple sclerosis (MS) and control (Ctrl) patients with idiopathic intracranial hypertension (IIH), across immune populations in blood and cerebrospinal fluid (CSF). For each comparison, the table reports sample size (*n*), mean percentage of cells expressing the gene in each group, the adjusted Wilcoxon rank-sum *P* value ($p_{adj}$), and the Hedges' *g* effect size with its 95% confidence interval (CI). *P* values were adjusted using Holm-Bonferroni correction for multiple comparisons within each gene and tissue (four comparisons per family). Positive Hedges' *g* values indicate a higher percentage of expressing cells in MS relative to IIH controls. Adjusted *P* values $< 0.05$ are shown in ***bold*** in the table. Hedges' *g* values with magnitude $\geq 0.8$ are also displayed in **bold** in the table.

| Protein Name | Amino Acid Sequence |
|---|---|
| WT IL-10 | RGQYSREDNNCTHFPVGQSHMLLELRTAFSQVKTFFQTKDQLDNILL<br>TDSLMQDFKGYLGCQALSEMIQFYLVEVMPQAEKHGPEIKEHLNSLG<br>EKLKTLRMRLRRCHRFLPCENKSKAVEQVKSDFNKLQDQGVYKAMN<br>EFDIFINYIEAYMMIKMKSHHHHHH |
| SA-IL-10 | EAHKSEIAHRYNDLGEQHFKGLVLIAFSQYLQKCSYDEHAKLVQEVT<br>DFAKTCVADESAANCDKSLHTLFGDKLCAIPNLRENYGELADCCTKQ<br>EPERNECFLQHKDDNPSLPPFERPEAEAMCTSFKENPTTFMGHYLH<br>EVARRHPYFYAPELLYYAEQYNEILTQCCAEADKESCLTPKLDGVKE<br>KALVSSVRQRMKCSSMQKFGERAFKAWAVARLSQTFPNADFAEITK<br>LATDLTKVNKECCHGDLLECADDRAELAKYMCENQATISSKLQTCCD<br>KPLLKKAHCLSEVEHDTMPADLPAIAADFVEDQEVCKNYAEAKDVFL<br>GTFLYEYSRRHPDYSVSLLLRLAKKYEATLEKCCAEANPPACYGTVL<br>AEFQPLVEEPKNLVKTNCDLYEKLGEYGFQNAILVRYTQKAPQVSTP<br>TLVEAARNLGRVGTKCCTLPEDQRLPCVEDYLSAILNRVCLLHEKTPV<br>SEHVTKCCSGSLVERRPCFSALTVDETYVPKEFKAETFTFHSDICTLP<br>EKEKQIKKQTALAELVKHKPKATAEQLKTVMDDFAQFLDTCCKAADK<br>DTCFSTEGPNLVTRCKDALAGGGSGGGSRGQYSREDNNCTHFPVG<br>QSHMLLELRTAFSQVKTFFQTKDQLDNILLTDSLMQDFKGYLGCQAL<br>SEMIQFYLVEVMPQAEKHGPEIKEHLNSLGEKLKTLRMRLRRCHRFL<br>PCENKSKAVEQVKSDFNKLQDQGVYKAMNEFDIFINYIEAYMMIKMK<br>SHHHHHH |

**Supplementary Table 2 | Amino acid sequences for murine WT IL-10 and SA-IL-10.** Sequences are shown from N- to C-terminus. WT IL-10 consists of murine IL-10 fused to a C-terminal hexahistidine tag for affinity purification. SA-IL-10 consists of murine serum albumin fused to murine IL-10 via a $(GGGS)_2$ flexible linker and includes a C-terminal hexahistidine tag for affinity purification. In both constructs, a single cysteine residue within murine IL-10 was substituted with tyrosine to reduce disulfide-mediated dimerization during expression and purification, as previously described.[71]

| Antibody | Source | Identifier |
|---|---|---|
| anti-CCR2, FITC (clone: SA203G11) | BioLegend | Cat. #: 150607<br>RRID #: AB_2616979 |
| anti-CCR7, PE-Cy7 (clone: 4B12) | BioLegend | Cat. #: 120123<br>RRID #: AB_2616687 |
| anti-CD8$\alpha$, BUV737 (clone: 53-6.7) | BD Biosciences | Cat. #: 612759<br>RRID #: AB_2870090 |
| anti-CD11b, BV650 (clone: M1/70) | BioLegend | Cat. #: 101239<br>RRID #: AB_11125575 |
| anti-CD11c, BV786 (clone: N418) | BioLegend | Cat. #: 117335<br>RRID #: AB_11219204 |
| anti-CD19, BV650 (clone: 6D5) | BioLegend | Cat. #: 115541<br>RRID #: AB_11204087 |
| anti-CD19, PerCP-Cy5.5 (clone: 6D5) | BioLegend | Cat. #: 115533<br>RRID #: AB_2259869 |
| anti-CD25, PerCP-Cy5.5 (clone: PC61) | BD Biosciences | Cat. #: 551071<br>RRID #: AB_394031 |
| anti-CD3, BV605 (clone: 17A2) | BioLegend | Cat. #: 100237<br>RRID #: AB_2562039 |
| anti-CD3, BUV615 (clone: 17A2) | BD Biosciences | Cat. #: 751418<br>RRID #: AB_2875417 |
| anti-CD4, BUV496 (clone: GK1.5) | BD Biosciences | Cat. #: 612952<br>RRID #: AB_2813886 |
| anti-CD40, BUV615 (clone: 3/23) | BD Biosciences | Cat. #: 751646<br>RRID #: AB_2875639 |
| anti-CD45, AF700 (clone: 30-F11) | BD Biosciences | Cat. #: 560510<br>RRID #: AB_1645208 |
| anti-CD45, BUV805 (clone: 30-F11) | BD Biosciences | Cat. #: 568336<br>RRID #: AB_3684191 |
| anti-CD86, BUV395 (clone: GL1) | BD Biosciences | Cat. #: 564199<br>RRID #: AB_2738664 |
| anti-CD206, AF700 (clone: C068C2) | BioLegend | Cat. #: 141733<br>RRID #: AB_2629636 |
| anti-CTLA-4, PerCP-Cy5.5 (clone: UC10-4B9) | BioLegend | Cat. #: 106315<br>RRID #: AB_2564473 |
| anti-CTLA-4, PE-Cy7 (clone: UC10-4B9) | BioLegend | Cat. #: 106313<br>RRID #: AB_2564237 |
| anti-F4/80, PE (clone: BM8) | BioLegend | Cat. #: 123109<br>RRID #: AB_893498 |
| anti-Foxp3, AF488 (clone: MF23) | BD Biosciences | Cat. #: 560403<br>RRID #: AB_1645192 |
| anti-GATA-3, BV421 (clone: L50-823) | BD Biosciences | Cat. #: 563349<br>RRID #: AB_2738152 |
| anti-ICOS, BV711 (clone: C398.4A) | BioLegend | Cat. #: 313547<br>RRID #: AB_2734288 |
| anti-I-A/I-E, BV421 (clone: M5/114.15.2) | BioLegend | Cat. #: 107631<br>RRID #: AB_10900075 |
| anti-PD-1, APC-Cy7 (clone: 29F.1A12) | BioLegend | Cat. #: 135223<br>RRID #: AB_2563522 |

| | | |
|---|---|---|
| anti-PD-L1, BV711 (clone: 10F.9G2) | BioLegend | Cat. #: 124319<br>RRID #: AB_2563619 |
| anti-RORγt, PE (clone: Q31-378) | BD Biosciences | Cat. #: 562607<br>RRID #: AB_11153137 |
| anti-Tbet, BV786 (clone: 4B10) | BioLegend | Cat. #: 644835<br>RRID #: AB_2721566 |
| anti-TIGIT, BV605 (clone: 1G9) | BioLegend | Cat. #: 142121<br>RRID #: AB_3083126 |
| $MOG_{38-49}$ Tetramer, APC | NIH Tetramer Core Facility | IEDB ID: 116101<br>GenBank ID: Q61885.1<br>Uniprot ID: Q61885 |

**Supplementary Table 3 | Antibodies used for flow cytometry.** Antibodies are listed with their target antigen, fluorophore conjugate, manufacturer, catalog number, and Research Resource Identifier (RRID).
